# Beyond a Linear Structure: The Tubular Organization of the Tripartite Attachment Complex and the Functional Role of TAC53

**DOI:** 10.1101/2025.05.14.653994

**Authors:** Clirim Jetishi, Salome Aeschlimann, Bernd Schimanski, Sandro Käser, Rachel Mullner, Silke Oeljeklaus, Bungo Akiyoshi, Bettina Warscheid, Falk Butter, André Schneider, Torsten Ochsenreiter

## Abstract

The Tripartite Attachment Complex (TAC) is essential for mitochondrial DNA (kDNA) segregation in *Trypanosoma brucei*, providing a physical link between the flagellar basal body and the mitochondrial genome. Although the TAC’s hierarchical assembly and linear organization have been extensively studied, much remains to be discovered regarding its complete architecture and composition – for instance, our identification of a new TAC component underscores these knowledge gaps. Here, we use a combination of proteomics, RNA interference (RNAi), and Ultrastructure Expansion Microscopy (U-ExM) to characterize the TAC at high resolution and identify a novel component, TAC53 (Tb927.2.6100). Depletion of TAC53 in both procyclic and bloodstream forms results in kDNA missegregation and loss, a characteristic feature of TAC dysfunction. TAC53 localizes to the kDNA in a cell cycle-dependent manner and represents the most kDNA-proximal TAC component identified to date. U-ExM reveals a previously unrecognized tubular architecture of the TAC, with two distinct TAC structures per kDNA disc, suggesting a mechanism for precise kDNA alignment and segregation. Moreover, immunoprecipitation and imaging analyses indicate that TAC53 interacts with known TAC-associated proteins HMG44, KAP68, and KAP3, forming a network at TAC–kDNA the interface. These findings redefine our understanding of TAC architecture and function and identify TAC53 as a key structural component anchoring the mitochondrial genome in *T. brucei*.

**Significance Statement:** This research identifies a new component (TAC53) of the tripartite attachment complex (TAC), a cellular machinery that anchors the mitochondrial DNA to a cytoskeletal structure, the basal body in *Trypanosoma brucei*. Using proteomics and high-resolution microscopy, we demonstrate that TAC53 is at the interface of the TAC and the mitochondrial DNA and likely the final piece in this structure. We also describe the overall tubular architecture of the TAC from the basal body to the mitochondrial matrix, and how the presence of two TAC structures per mitochondrial genome can explain the parasite’s ability to maintain its mitochondrial DNA accurately. In summary, we present a new component and the architecture of the currently best understood mitochondrial DNA segregation mechanism in biology.

## Introduction

*Trypanosoma brucei* is a single-celled eukaryotic parasite. Major discoveries made in trypanosomes like GPI anchors, trans splicing, RNA editing and acidocalcisomes were later also found to be important features in other eukaryotes (1–4). Trypanosomes have served as a model system to study mitochondrial biogenesis in an organism evolutionary distinct from the mainstream systems like yeast and human cells. Unlike yeast and mammals, *T. brucei* cells harbor only one single mitochondrion, with a single nucleoid that is named kinetoplast and contains the mitochondrial DNA (kinetoplast DNA, kDNA) (5–8).

The kDNA consists of two different genetic components, the maxi- and the minicircles, which are topologically interlocked and form a dense disc-shaped structure. While the maxicircles encode for ribosomal RNAs and proteins required in the respiratory chain and the mitochondrial ribosome, the minicircles encode for guide RNAs (gRNAs) which are required for editing the maxicircle encoded transcripts (9–11). The replication of trypanosomal kDNA, unlike other eukaryotic mitochondrial genomes, is coordinated with the nuclear cell cycle and the segregation of replicated kDNA discs is completed before mitosis (12, 13). The single-unit nature of the kDNA requires a precise mechanism to ensure that both daughter cells inherit one kDNA disc. This is facilitated by a stable connection of the kDNA disc with the flagellum’s basal body. This linkage ensures the segregation of both the old and new flagellum along with the replicated kDNA (14).

The Tripartite Attachment Complex (TAC) is the structure that physically connects the basal body to the kDNA disc and positions the kDNA at the posterior end of the cell (15). The TAC consists of three distinct subdomains: i) the exclusion zone filaments (EZF) between the basal body of the flagellum and the outer mitochondrial membrane (OMM), ii) the differentiated membranes (DM) where the inner mitochondrial membrane (IMM) is closely opposed to the OMM and iii) the unilateral filaments (ULF) that connect the IMM to the kDNA disc (6, 16, 17). Although mitochondrial inheritance in other organisms involves cytoskeletal elements like actin and microtubules, a stable, permanent connection between mitochondrial nucleoids and the cytoskeleton has only been described in *T. brucei* (17–20). Therefore, the TAC provides a direct physical link between the mitochondrial DNA and the cytoskeleton, making it the only known example of such a permanent DNA-cytoskeleton connection.

So far eight TAC proteins have been described in *T. brucei* (21). The component closest to the BB and potentially directly connected to it, is the approximately 880 kDa protein p197 (22–25). It has been shown that p197 filaments determine the distance between the BB and the OMM, where it directly binds TAC65 (24). TAC65 is a peripherally OMM associated protein that interacts with the OMM integrated pATOM36. Interestingly, pATOM36 not only operates as a TAC protein, but also functions in biogenesis of a subset of OMM proteins. Therefore, pATOM36 is located throughout the entire OMM (26, 27). TAC40, TAC42 and TAC60 form a complex in the OMM. TAC40 and TAC42 are β-barrel proteins, whereas TAC60 is a -helical protein with two transmembrane domains (TMDs) (28, 29). Both the N- and C-termini of TAC60 are located in the cytoplasm, resulting in the formation of a loop in the inter membrane space (IMS) (28) For the IMM region of the TAC, only one protein is described, p166. Being the first TAC protein to be identified in 2008 (30), it was recently shown to have a mitochondrial targeting sequence (MTS) and being integrated into the IMM with single C-terminal TMD. The very C-terminus of p166 reaches into the IMS and interacts with the loop formed by TAC60 (31, 32). The N-terminus of p166 directly interacts with TAC102, a protein of the ULFs (33). TAC102 was identified as the most kDNA-proximal TAC component, however, direct kDNA binding could not be assigned (23, 34). In addition to the TAC proteins, two non-TAC proteins, HMG44 and KAP68, were suggested to interact with TAC102. These proteins were shown to interact with each other in vitro, and both their depletion results in kDNA loss. Since HMG44 and KAP68 are located closer to the kDNA than TAC102, it was proposed that they anchor the TAC to the kDNA (35).

A shared feature of all TAC proteins is that their depletion leads to a missegregation phenotype, where the replicated kDNA will stay attached to the “old” TAC, whereas the “new” TAC emanating from the newly assembled pro-basal body only has fractions or no kDNA. Ultimately this leads to kDNA loss in the daughter cells, while the mother cell retains the over-replicated kinetoplast (28). Additionally, TAC proteins localize to the TAC regions in intact cells as well as isolated flagella (17) and they are dispensable in an engineered cell line of the bloodstream form of *T. brucei* (γL262P), which can survive without the kDNA (36).

Since the TAC assembles in a hierarchical manner, the depletion of upstream components (basal body proximal) will lead to mislocalization and depletion of downstream components (kDNA proximal proteins) (23). We made use of the TAC hierarchy, and performed TAC-depletome experiments with isolated flagella on p197 and TAC42, identifying a new TAC protein, TAC53.

Ultrastructure Expansion Microscopy (U-ExM) with improved structural integrity and better preservation of ultrastructural details (37) were successfully established in the protozoan model systems like *Trypanosoma brucei*, *Leishmania major*, *Toxoplasma gondii, Giardia lamblia* and several *Plasmodium* species, providing insights into their unique cellular architecture (38–44). In *T. brucei,* U-ExM not only allowed intermolecular information, but also allowed the localization of two distinct domains within the same protein (24).

In this study, we employed U-ExM to investigate the TAC, which has been traditionally observed from a lateral perspective. However, for the first time, we also analyzed it from the axial perspective, revealing that the TAC forms a tubular structure, indicated by the ring-like formation seen from the axial perspective. This finding provides new insights into the structural organization of the TAC and suggests a more complex architecture than previously thought.

## Results

### TAC depletomes reflect TAC biology

To identify new TAC components, we performed TAC depletomic assays as previously described. In brief, the kDNA remains attached to the basal body via the TAC during isolation of the flagellum (15, 27, 35). We have also shown that the TAC assembles in a hierarchical fashion such that if you deplete a component of the TAC only the kDNA proximal part of the TAC will be lost while the basal body proximal part remains (23). We thus can compare the composition of the TAC and its associated proteins from cells depleted of individual TAC components by quantitative mass spectrometry (MS). For this, stable isotope labelling by amino acids in cell culture (SILAC) in combination with quantitative MS has been performed using isolated flagella (23, 35). To avoid the detection of proteins like replication factors that are only connected to the TAC via the kDNA, a DNase digestion was performed before samples were analyzed by MS.

### p197 depletomics

In the depletomic experiment targeting p197 (Tb927.10.15750, Tb11.v5.0394) we detected 1665 proteins, 43 of which were depleted >1.5 fold including all known TAC components (Figure 1A, Supplementary Table S1). Other proteins that were affected by the depletion of p197 include a basal body protein, several components of the oxidative phosphorylation chain as well as proteins of unknown function. From the latter we selected seven candidates for further analysis based on their predicted localization to the mitochondrion or the basal body according to *TrypTag* (*45–47*). To test their potential function in the TAC we targeted the seven candidates by RNAi in the γL262P cell line (Supplementary Figure S1) (36). We monitored growth as well as kDNA replication and segregation using DAPI staining. Depletion of only one candidate (Tb927.5.2970) showed loss and over-replication of the kDNA as well as a growth arrest. While the over-replication might hint towards a function in the TAC the rapid growth defect in the γL262P cell line indicates that the protein is involved in basal body biogenesis rather than mitochondrial genome inheritance. In summary none of the tested candidates fulfill the criteria of a TAC component.

**Figure 1.**
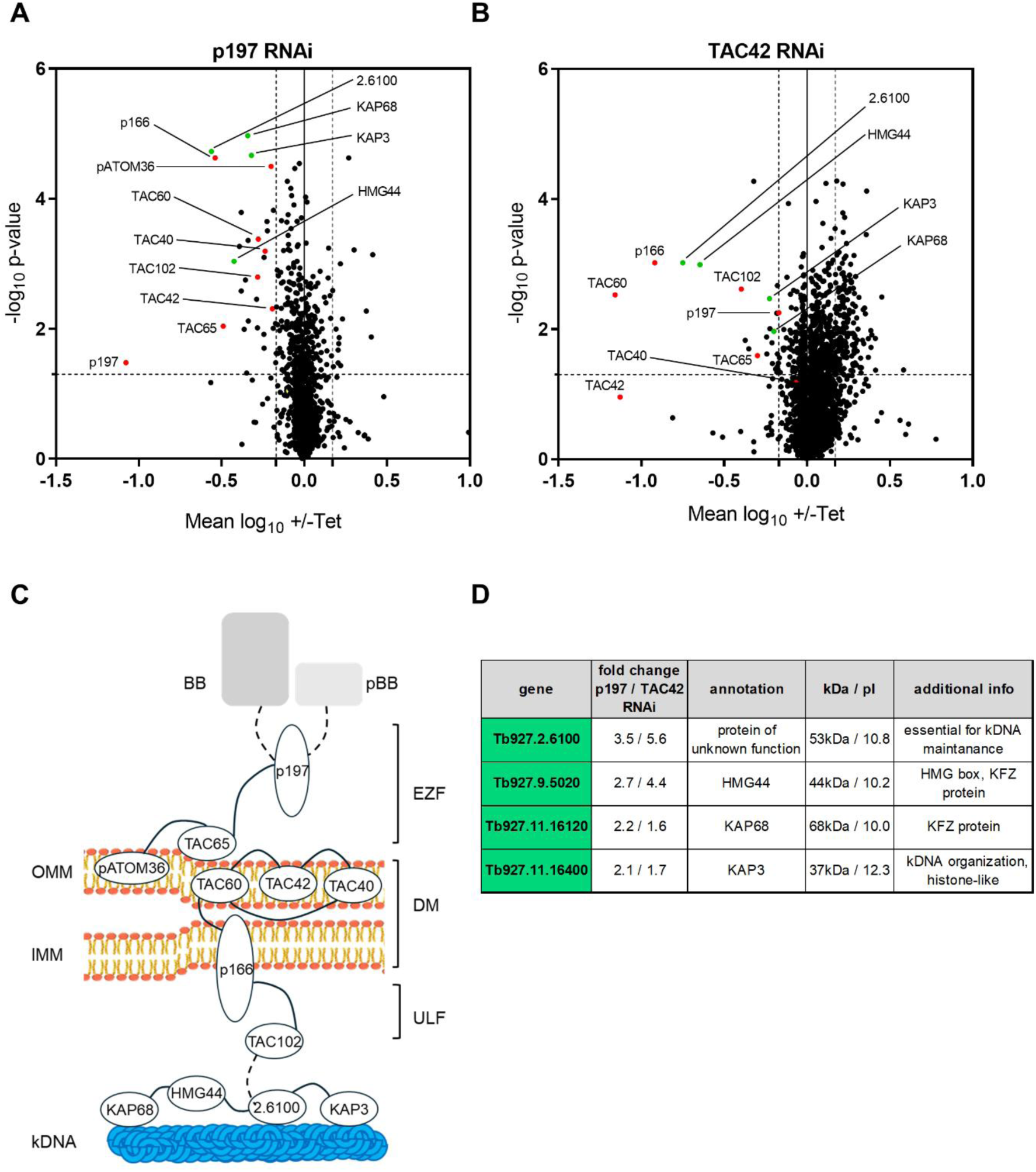
**Depletomic analysis targeting the TAC proteins p197 and TAC42.** AB) Volcano plots showing the mean log10 ratios (induced/uninduced) of normalized protein abundance from four (A) or three (B) biological replicates plotted against the - log10 p-values (two-sided Student’s t-test). The dashed vertical lines refer to the 1.5-fold change threshold and the dashed horizontal lines mark the significance line (p<0.05). Complete lists of all proteins that were depleted more than 1.5 times are shown in the supplementary tables S1 and S2. Known TAC components are depicted in red; proteins without a TAC assignment that were depleted >1.5 with a p-value < 0.05 present in both datasets (A and B) are depicted in green. (C) Schematic representation of the Tripartite Attachment Complex TAC. TAC - and TAC-associated proteins are shown. Connecting lines indicate known interactions, dashed line possible interactions. BB, basal body; pBB, pro-basal body; OMM, outer mitochondrial membrane; IMM, inner mitochondrial membrane; EZF, exclusion zone filaments; DM, differentiated membranes; ULF, unilateral filaments. (D) Table of proteins without TAC assignment that were depleted >1.5 with a p-value < 0.05 in both datasets.

### TAC42 depletomics

In the TAC42 (Tb927.7.3060) depletomic experiment we detected 2575 proteins, 52 of which were depleted >1.5 fold including five known TAC components (Figure 1B, Supplementary Table S2). Other proteins that were affected by the depletion of TAC42 include a basal body protein, several proteins involved in flagellar biogenesis as well as proteins of unknown function. From the latter we again selected seven candidates for further analysis based on their predicted localization to the mitochondrion according to *TrypTag* (*45–47*). We tagged the proteins and defined their localization *in situ* and selected three candidates that localize to the kDNA for functional studies by RNAi (Tb927.10.8980, Tb927.1.2730, Tb927.8.3160) (Supplementary Figure S2). Targeting these candidates by RNAi leads to a growth defect, but the kDNA is not affected. Again, none of the tested candidates fulfill the criteria of a TAC component.

We also noticed that aside from the known TAC components four proteins were significantly depleted >1.5 fold in both data sets (Tb927.2.6100; Tb927.9.5020, HMG44; Tb927.11.16120, KAP68; Tb927.11.16400, KAP3). These four proteins have previously been studied in detail and have been proposed to be involved in functions other than the TAC. HMG44 and KAP68 are part of a distinct complex connecting the kDNA to the TAC (35). KAP3 is a histone H1 like protein involved in kDNA organization and Tb927.2.6100 was shown to be involved in kDNA maintenance (48–50). Interestingly, all four proteins were significantly depleted >2-fold in a RNAi cell line targeting TAC102, the protein previously thought to be the closest TAC component to the kDNA (35). The above experiments suggest that all four proteins are positioned closer to the kDNA within the TAC hierarchy.

Since Tb927.2.6100 was repeatedly identified in our screens for new TAC components we decided to further characterize this protein (23, 33, 35).

### Tb927.2.6100 (TAC53) is an essential protein of the TAC

The 53 kDa Tb927.2.6100 is a protein essential for normal growth with a pI of 10.77 and a predicted N-terminal mitochondrial targeting signal. It has putative homologs in most trypanosomatids including *Leishmania* and *Paratrypanosoma* (Supplementary Figure S3). Interestingly, minor groove DNA-binding motifs such as AT hooks and SPKK motifs are present in some homologs, suggesting a putative role in interaction with the kDNA (51, 52). We generated an inducible RNAi cell line targeting Tb927.2.6100. Consistent with previous findings, depletion of Tb927.2.6100 resulted in a slow growth phenotype three to four days post-induction (50) (Figure 2A). Additionally, cells depleted of Tb927.2.6100 show a kDNA loss phenotype with less than 20 % of cells harboring a kinetoplast three days after RNAi induction in PCF (Figure 2A).

**Figure 2.**
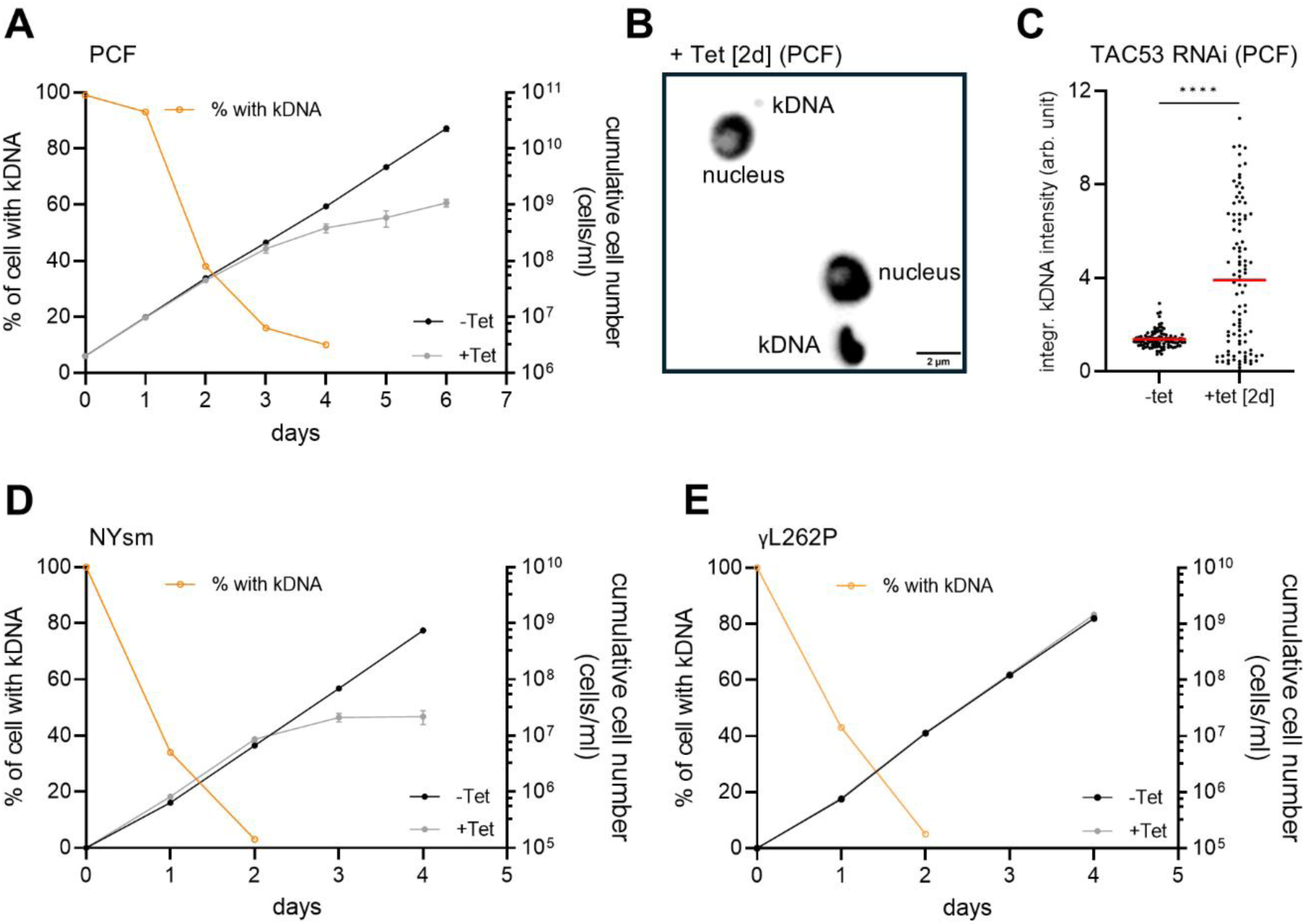
**Tb927.6100 encodes for the essential TAC component TAC53.** A) Growth curve of the procyclic TAC53 RNAi cell line. The orange line depicts the percentage of cells with kDNA, while the black line shows the growth curve of uninduced cells, and the grey line shows RNAi-induced cells (induced with tetracycline). B) Fluorescence DAPI stain of two representative cells after TAC53 RNAi [2 days induced]; The cell on the top shows a tiny kDNA, the bottom cell shows an over-replicated kDNA. C) Integrated kDNA intensities were measured in uninduced and RNAi-induced cells [2 days]. The red line marks the mean. D, E) Growth curves and loss of kDNA of TAC53 RNAi in depicted bloodstream form host cell lines. Color code of graph as in (A).

Using fluorescence microscopy, we also observed kDNA missegregation, which was not reported in the previous study but is typical for the loss of function phenotype of TAC proteins (50) (Fig 2 B). To quantify the observations, we measured integrated kDNA intensities in cells where Tb927.2.6100 was depleted by RNAi for two days and compared it to uninduced cells (Figure 2C).

To further test if Tb927.2.6100 functions as a TAC protein, we repeated the RNAi experiments in wildtype NYsm bloodstream forms and in the γL262P bloodstream form cell lines (17, 36). In both cell lines depletion of Tb927.2.6100 leads to loss of kDNA in more than 90% of the cells.

While this leads to a severe growth defect in the NYsm, the γL262P cell line growth is not affected (Fig 2 D and E).

Based on these findings, we conclude that Tb927.2.6100 is an essential TAC protein and will be named TAC53.

### TAC53 localizes the kDNA in a cell-cycle dependent manner

To investigate the subcellular localization of TAC53, a polyclonal antibody was generated, and its specificity was validated by RNAi in conjunction with immunofluorescence microscopy and Western blot analysis (Supplementary Figure S4). Subsequent immunofluorescence assays utilizing this antibody, in combination with DAPI staining, confirmed the subcellular localization of TAC53 at the kDNA (Figure 3A). Quantification of the integrated fluorescence signals, normalized to the kDNA, showed that TAC53 signals vary during the cell cycle. TAC53 is significantly enriched in cells undergoing kDNA division (1kdiv1N) and in cells with two kinetoplasts and one nucleus (2k1N), compared to cells with a single kinetoplast and nucleus (1k1N) or two kinetoplasts and two nuclei (2k2N) (Figure 3B). In contrast, TAC102 levels remained unchanged throughout the cell cycle (Figure 3C).

**Figure 3.**
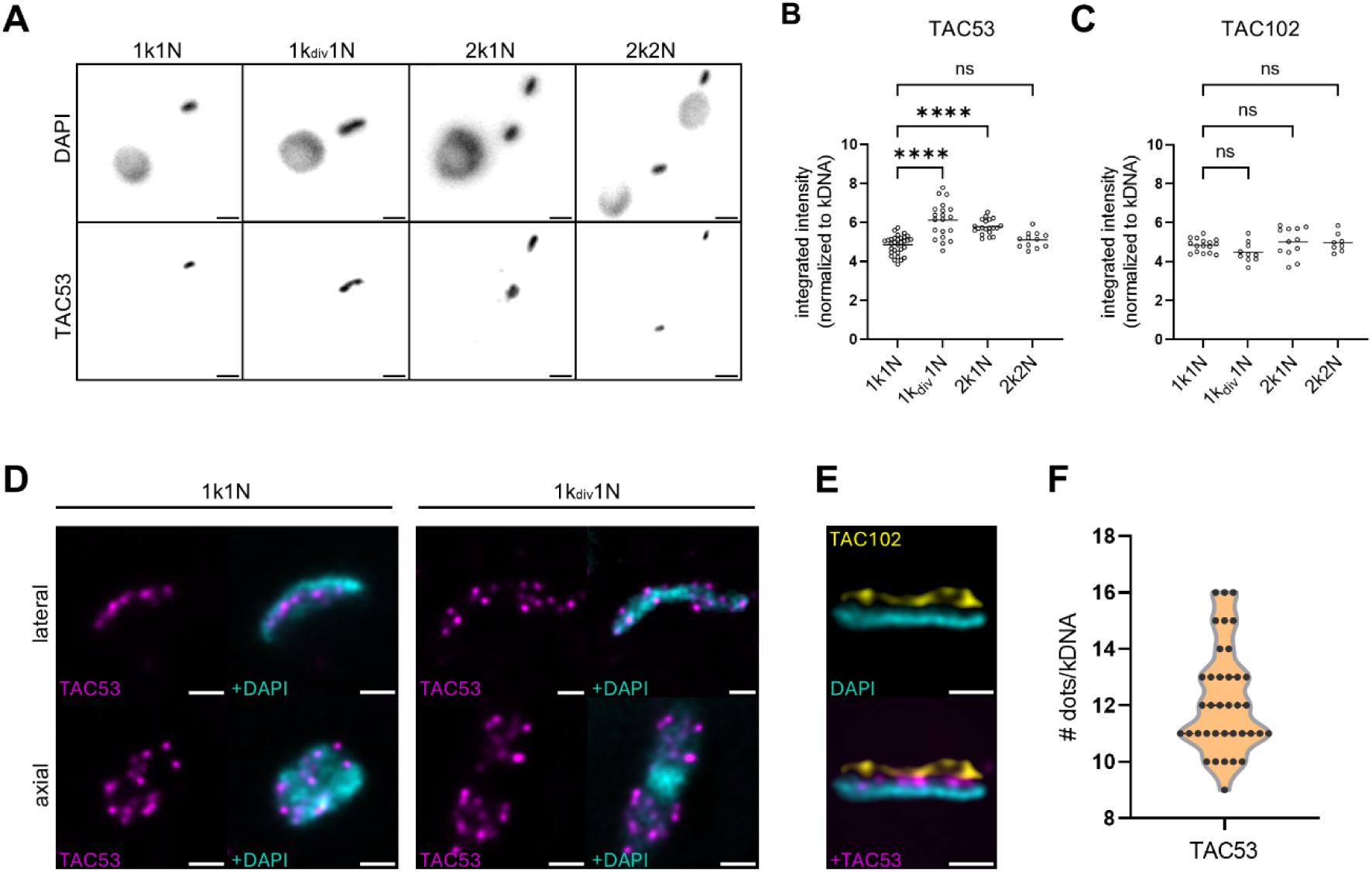
**TAC53 localizes at the kDNA in a cell-cycle specific manner.** A) Immunofluorescence images of wildtype cells in different cell cycle stages stained with the TAC53 antibody. The upper lane shows DAPI signal, the lower lane shows signal for TAC53. B) Quantification of integrated signal intensities for TAC53 in different cell cycle stages, normalized to the kDNA. C) Quantification of integrated signal intensities for TAC102 in different cell cycle stages, normalized to the kDNA. D) U-ExM of TAC53 signal in 1k1N cells (left) and 2k1N cell (right) in magenta. kDNA is depicted in cyan. E) U-ExM co-staining of TAC102 (yellow) and TAC53 (magenta), with kDNA in cyan. Scale bars in D) and E) represent 1 µm (not adjusted for Expansion). F) Quantification of the punctate signals seen in U-ExM images for TAC53 per kDNA, n=37 cells.

To further resolve the precise spatial distribution of TAC53, U-ExM was employed, a high-resolution imaging technique that enhances spatial resolution through physical expansion of the specimen (37). In 1k1N cells TAC53 was detected as a punctate signal distributed throughout the entire kDNA disc, however, in 1kdiv1N cells, TAC53 signal appears on both kDNA lobes with a similar pattern as in 1k1N with little to no presence in center of the kinetoplast. (Figure 3D).

Additionally, co-staining with a monoclonal antibody against TAC102 revealed that TAC53 is positioned more proximally to the kDNA compared to TAC102 (Figure 3E, right). This observation, along with findings from depletomic assays, indicates that TAC53 represents the TAC component most closely associated with the kDNA. In the expanded cells, we additionally quantified the punctate signals from TAC53 staining as shown in Figure 3F. Our analysis, based on 37 measured kDNA lobes, indicated that the majority of kDNA lobes exhibited predominantly 11 foci, with a range of nine to 16 foci observed.

### TAC53 incorporation into the nascent TAC depends on the kDNA

Since the TAC53 signal fluctuates throughout the cell cycle and is most abundant in segregating kDNAs, we sought to determine whether the kDNA is required for the proper localization of TAC53. To explore this, we utilized the γL262P cell line, which is particularly advantageous for studying the TAC because it permits a reversible disruption of TAC biogenesis. Upon tetracycline-inducible RNAi against a TAC component, the TAC structure is disrupted, leading to loss of kDNA. Once the kDNA has been eliminated, the TAC component targeted by RNAi is re-expressed by removal of tetracycline from the media, allowing TAC reassembly to occur in the absence of kDNA (35).

Since the TAC assembles hierarchically from the basal body to the kinetoplast, depletion of basal body-proximal TAC components leads to the loss of more distal components. Consequently, p197 RNAi depletes all TAC proteins, including TAC102 and TAC53. Immunofluorescence microscopy revealed that five days after p197 RNAi induction, only approximately 10% of the cells retained a detectable TAC102 signal, while no cells exhibited TAC53 signal (Figure 4A, middle lane). In contrast, in uninduced cells, both proteins were detected in over 95% of the population (Figure 4A, upper lane). Upon p197 re-expression, nearly 90% of cells successfully reassembled TAC102 into the TAC, consistent with previous findings (35), whereas only 20% of cells restored TAC53 incorporation at this time point (Figure 4A, lower lane).

**Figure 4.**
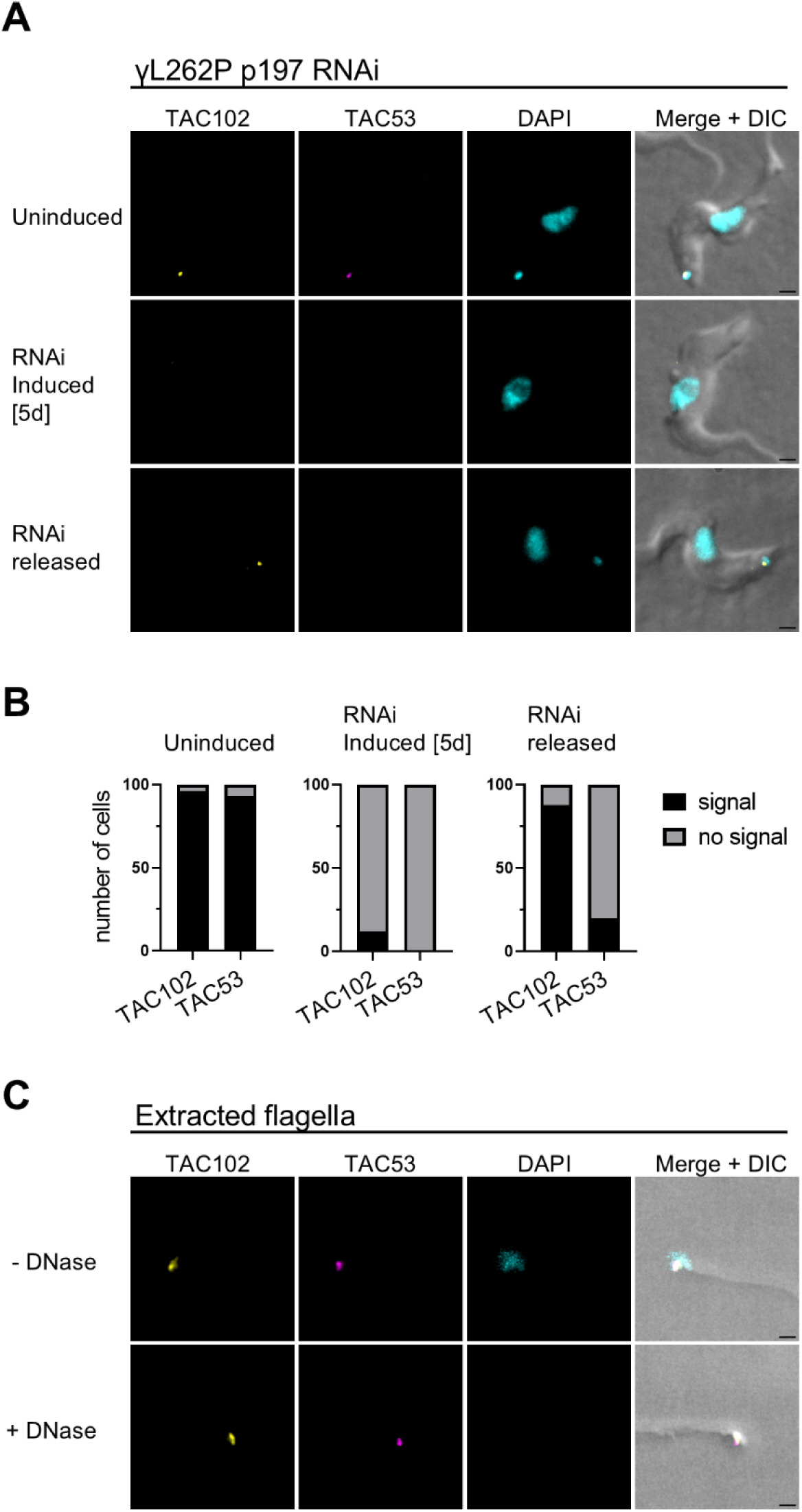
**Dynamics of TAC53 and TAC102 During TAC Assembly and in Isolated Flagella.** A) Immunofluorescence analysis of TAC53 and TAC102 relative to the kDNA during TAC depletion and reassembly in the γL262P-p197 RNAi cell line. The top row shows uninduced cells, the middle row shows cells after five days of RNAi induction, and the bottom row shows cells following re-expression of p197. TAC102 is shown in yellow, TAC53 in magenta, and DNA in cyan. The scale bar is 1µm. B) Quantification of signal intensities shown in (A). C) Immunofluorescence analysis of TAC53 and TAC102 in extracted flagella, either untreated (top row) or treated with DNase-I (bottom row). Color coding as in (A). The scale bar is 1µm.

To assess whether TAC53, once assembled into the TAC, still requires kDNA for its stable association with the structure, we conducted immunofluorescence microscopy on DNase I-treated isolated flagella. The results demonstrated, in agreement with the SILAC pulldown experiments (Fig. 1), that TAC53 remains attached to the isolated flagella (Figure 4C), similar to previous findings for TAC102 (35). This suggests that once TAC53 is incorporated into the TAC, its localization is no longer dependent on the presence of kDNA.

### Immunoprecipitation and U-ExM reveal protein interactions and localization

TAC53 appears to be the kDNA most proximal component of the TAC structure and thus is in close proximity to the recently described HMG44 and KAP68 (35). To test if HMG44 and KAP68 form a complex with TAC53 *in situ*, we used immunoprecipitation assays with cell lines expressing HA- and myc-tagged KAP68 and HMG44, respectively. Consequently, mitochondrial enriched fractions were sonicated, and the resulting soluble fractions were subjected to immunoprecipitations using anti-myc or anti-HA beads. In the KAP68 immunoprecipitation we identified 27 significantly enriched proteins (enrichment >2 fold). Among those 16 have a mitochondrial annotation and seven are predicted to be localized to the kDNA including HMG44 (Figure 5A, Supplementary Table S3). In the HMG44 immunoprecipitation experiment we identified 11 significantly enriched proteins (enrichment >2 fold). Among these proteins, five have a predicted mitochondrial localization, including KAP68 and TAC53 (Figure 5B, Supplementary Table S4).

**Figure 5.**
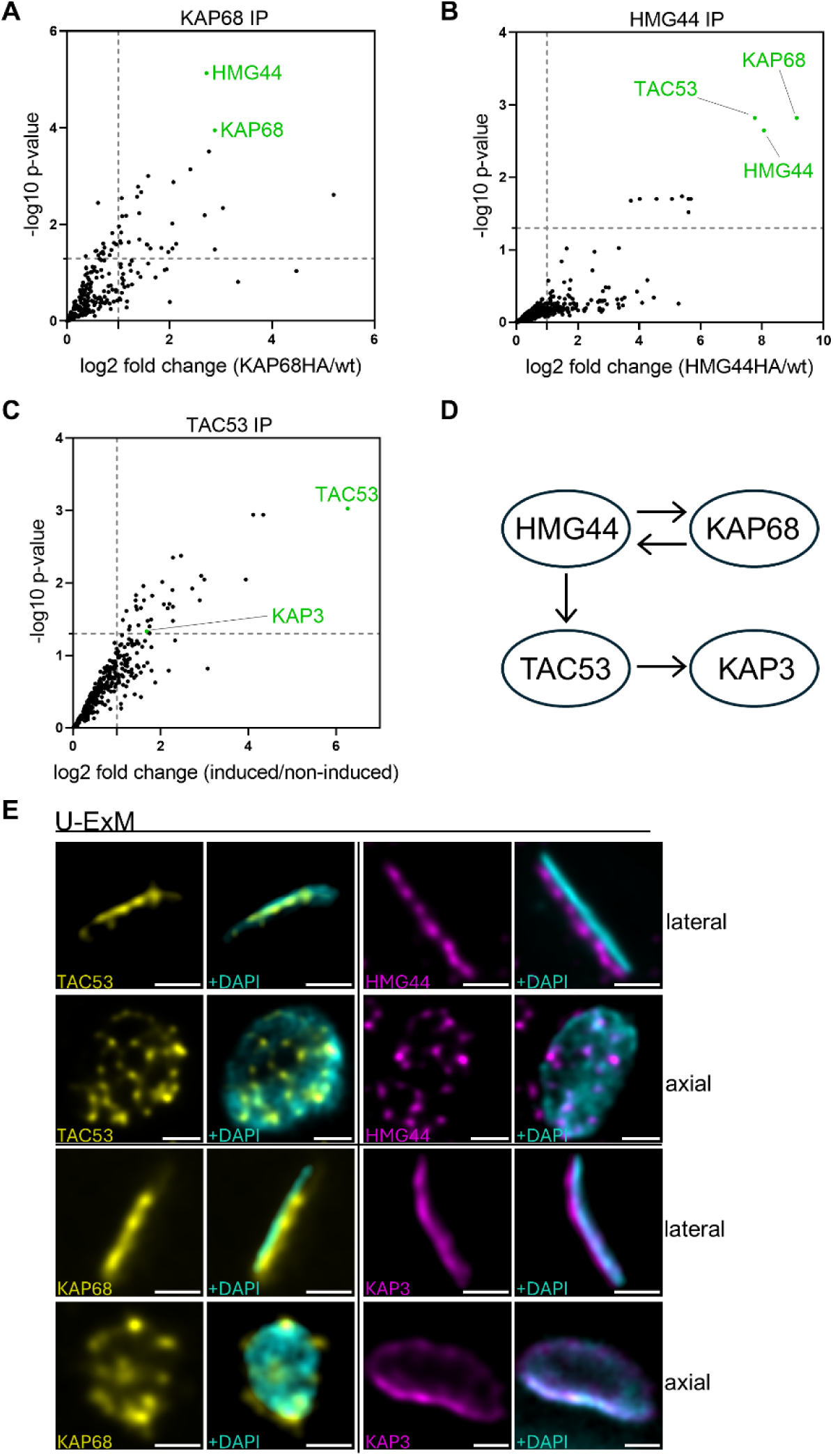
**Immunoprecipitation of C-terminally tagged HMG44, TAC53 and KAP68 from sonicated digitonin-enriched organelles.** A) Volcano plots showing the log2 fold changes (KAP68 /wt) of protein abundance plotted against the -log10 p-values from four technical replicates. The dashed vertical lines refer to the 2-fold change threshold and the dashed horizontal lines mark the significance line (p<0.05). B) Volcano plots showing the log2 fold changes (HMG44 /wt) of protein abundance plotted against the -log10 p-values from three technical replicates. The dashed vertical lines refer to the 2-fold change threshold and the dashed horizontal lines mark the significance line (p<0.05). C) Volcano plots showing the log2 fold changes (TAC53-induced/TAC53-uninduced) of protein abundance plotted against the -log10 p-values from three technical replicates. The dashed vertical lines refer to the 2-fold change threshold and the dashed horizontal lines mark the significance line (p<0.05). D) Schematic representation of the interactions based on the immunoprecipitation experiments. E) U-ExM images of TAC53, HMG44, KAP68 and KAP3 shown from a lateral and an axial view. TAC53 and KAP68 are shown in yellow, HMG44 and KAP3 are shown in magenta, kDNA is shown in cyan. The scale bar is 1µm (not adjusted for Expansion).

To identify TAC53 interaction partners, we used a cell line overexpressing TAC53-HA upon tetracycline induction, in the background of a 3’ UTR RNAi targeting endogenous TAC53. This construct partially rescues the 3’ UTR RNAi phenotype, showing reduced and non-significant kDNA missegregation, and a delayed growth phenotype compared to the 3’ UTR RNAi condition (Supplementary Figure S5 A-D). While the TAC53-HA construct localizes similarly to the endogenous protein, it also induces a growth phenotype when overexpressed in the 29-13 host cell line (Supplementary Figure S5 E, F). The observed growth phenotype may result from the HA-tag at the C-terminus of the TAC53-HA construct, which potentially disrupts the protein’s normal function. Additionally, the excessive expression of TAC53, which is regulated cell cycle dependent, could lead to a disruption in its developmental control due to elevated protein levels.

In the immunoprecipitation a total of 35 proteins were significantly enriched including the bait TAC53 (Figure 5C, Supplementary Table S5). Although many of the enriched proteins are annotated with cytoplasmic or nuclear localization, two proteins, in addition to TAC53, are predicted to localize to the kinetoplast. These are KAP3 (see Figure 1D) and the kDNA-associated protein Tb927.10.8980. However, the latter was identified by MS with a p-value > 0.05 (Supplementary Figure S2, Supplementary Table S5).

Additionally, we performed U-ExM for a better understanding of the relative orientation of the four proteins TAC53, HMG44, KAP68 and KAP3 at the interface of the TAC and the kDNA. From the lateral perspective, all observed proteins are closely associated with the kDNA. However, from the axial view, HMG44 and KAP68 exhibit a distribution pattern similar to TAC53, forming distinct punctate structures throughout the kDNA. In contrast, KAP3 seems to laterally encircle the kDNA. These findings indicate that the four proteins establish an interaction network, as evidenced by immunoprecipitation assays and U-ExM.

### U-ExM of the TAC reveals a tube-like structure

To characterize the overall structure of the TAC we pursued U-ExM. Using double tagged cell lines in combination with available antibodies we imaged the overall architecture of the TAC from two main orientations relative to the kDNA disc, lateral and axial.

The TAC is organized in three regions: the EZF represented by p197, the DM represented by TAC65, pATOM36, TAC40, TAC42, TAC60 and p166 which transitions into the ULF that also contain TAC102 and potentially TAC53 (see Figure 1C).

The C-termini of p197 form a circular structure with a mean diameter of 0.89 ± 0.08 µm and are close to the proximal end of the pro- and mature basal bodies, which have a mean diameter of 0.85 ± 0.07 µm (expanded cells, Figure 6 A, I). The N-termini of p197 also form a circular structure with a mean diameter of 1.07 ± 0.14 µm and are close to the OMM (Figure 6A). Thus, both the mature basal body and the pro-basal body each have one distinct TAC structure per kDNA prior to replication of the disc (Figure 6A).

**Figure 6.**
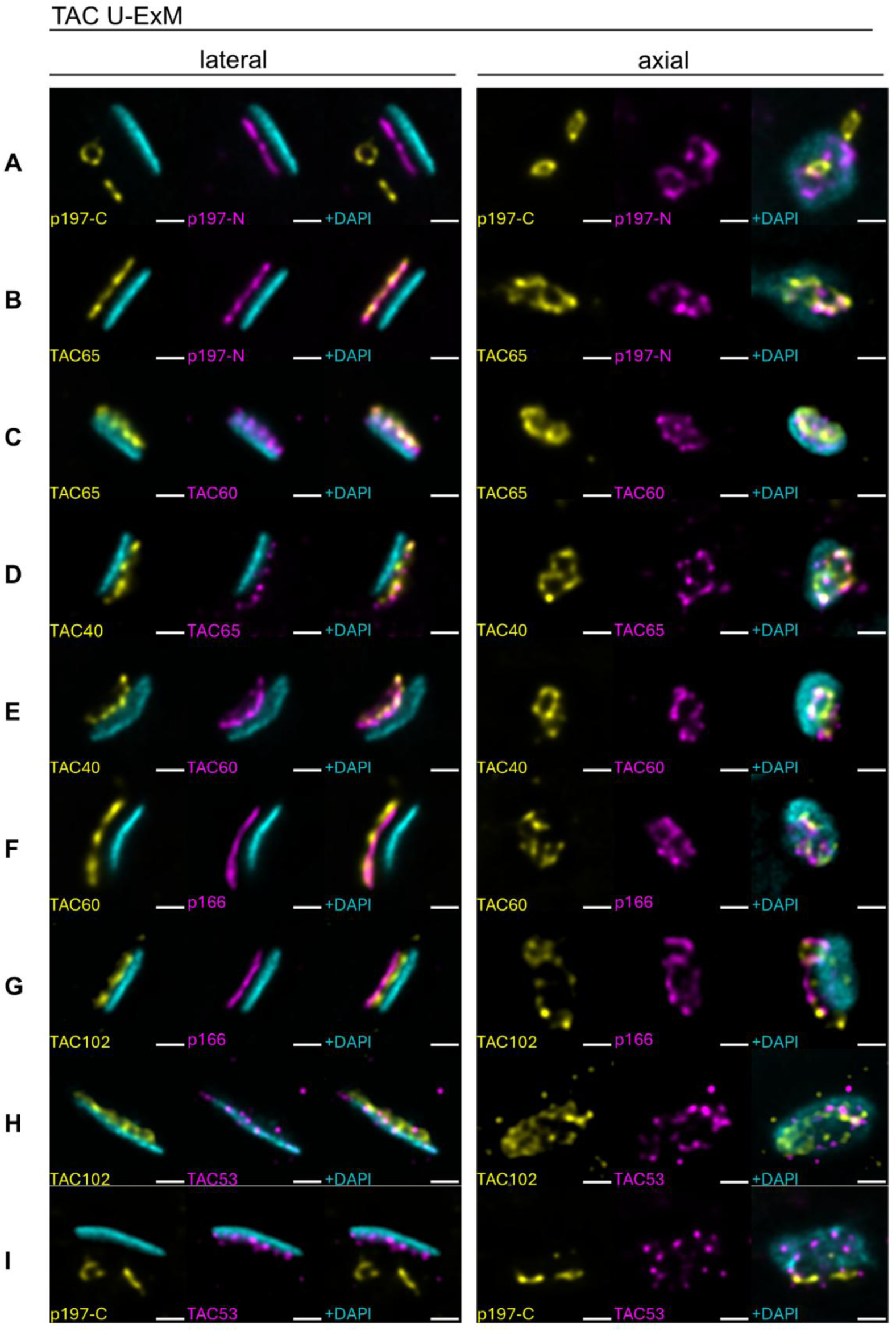
**Representative U-ExM images of TAC proteins in double-stained cells** Rows A–I show different TAC protein combinations. Each row corresponds to one protein pair, with a lateral view (left) and an axial view (right). Within each column, color coding remains consistent across all panels. Proteins imaged are indicated in the respective panels. In the lateral view, the first column displays the first protein in yellow and DNA in cyan, the second column shows the second protein in magenta with the same DNA signal, and the third column presents the merged image. In the axial view, the first and second columns show only the respective proteins (yellow or magenta), while the third column again shows the merge of both proteins. The scale bar is 1µm (not adjusted for Expansion).

In the DM, TAC65 follows the circular architecture of p197, with two distinct rings visible for each kDNA (Figure 6 B-C). Similarly, other OMM components of the TAC including TAC40 and TAC60 show distinct circular structures (Figure 6 C-F). Also, the IMM and ULF protein p166 exhibits a circular architecture (Figure 6 F-G). The diameters of the TAC structure seem to enlarge and diffuse from the basal body towards p166 (Figure 6J). While the TAC diameter marked by the C-terminus of p197 (0.89 ± 0.08 µm, expanded cells) is still very close to the basal body diameter (0.85 ± 0.07 µm, expanded cells), it is about 30% larger for the TAC components of the OMM (e.g., TAC40, 1.21± 0.17 µm, expanded cells, Figure 6J).

While p166 in the ULF still forms a circular structure, the signal for TAC102 which directly associates with p166 is more diffuse (Figure 6 G-H). The newly discovered TAC component TAC53 is visible as individual puncta with no specific arrangement (Figure 6 H-I).

We additionally imaged cells just prior to kDNA segregation when the kDNA is essentially fully replicated and used a combination of tagged p197 (both termini) and TAC53 (Supplementary Figure S6) to visualize the positions of these proteins. For the C-terminus of p197 we detected four distinct TAC structures with a clear separation, while the N-terminus of p197 only shows this degree of separation between the basal body pairs, (and less between pro- and mature basal bodies). TAC53 on the other hand keeps the punctate signal but is not present between the two segregating kDNA discs.

These findings support the presence of two distinct TAC structures prior to kDNA replication, forming a tube-like configuration.

## Discussion

This study advances our understanding of the architecture and molecular composition of the tripartite attachment complex in *T. brucei*, offering significant insights into the mechanisms underlying mitochondrial genome segregation in this eukaryotic model organism. By employing a multidisciplinary approach integrating quantitative proteomics, RNA interference, and ultrastructure expansion microscopy, we delineated the structural organization of the TAC and identified TAC53 as a previously uncharacterized component of the complex.

Application of U-ExM revealed that, from an axial viewpoint, the TAC exhibits a tubular morphology reminiscent of a truncated cone, indicative of a more intricate three-dimensional organization than previously appreciated. Starting from the C-terminal region of p197, a protein localized within the exclusion zone filaments, the tubular structure conforms closely to the barrel-shaped architecture of the basal body. Progressing toward the N-terminal region of p197 and approaching the outer mitochondrial membrane, the TAC displays a gradual increase in diameter (Supplementary Figure S7). This morphological gradient suggests an expansion along the basal body–kDNA axis, forming a structure resembling a truncated cone. At the outer mitochondrial membrane, proteins including TAC65, TAC40, and TAC60 are arranged in a circular manner that align with the N-terminus of p197 (Figure 6). This structural expansion is especially prominent in TAC40, which presents a diameter approximately 30% greater than that of the basal body. Such an enlargement may be necessary for the TAC to establish a robust physical interface with the more voluminous kDNA structure. While the inner mitochondrial membrane protein p166 maintains a circular geometry this structural definition diminishes in the unilateral filaments as evidenced by the diffuse U-ExM signal of TAC102 and the punctate pattern of TAC53, the newly identified TAC component.

Building on a previously proposed model of kDNA replication – in which each kDNA network comprises a diploid set of minicircles which is distributed across each lobe of the kDNA network – our findings provide compelling morphological evidence in support of this model (53). Specifically, U-ExM analysis demonstrated that in non-dividing cells each kDNA network is associated with two TAC structures: one connected to the mature basal body and another to the pro-basal body (Supplementary Figure S6E). This architectural configuration may constitute the structural basis for spatial segregation of the largely diploid mitochondrial genome within the kDNA disc. Furthermore, such spatial organization may also underlie the formation of the two replication foci known as the antipodal sites.

Given the TAC’s hierarchical organization – wherein proximal components near the basal body are prerequisite for the assembly of more distal elements – we employed a SILAC-based depletomic strategy to systematically identify novel TAC components. This approach, which enables unbiased detection of protein dependencies, successfully recapitulated the identification of several previously characterized TAC components (Figure 1A, B). Through extensive validation of novel candidate proteins derived from two independent screening datasets, we identified TAC53 as a bona fide and likely final structural component of the TAC. RNAi-mediated knockdown of TAC53 resulted in pronounced TAC missegregation phenotypes, including loss of kDNA and kDNA over-replication in both the procyclic and bloodstream life cycle stages of *T. brucei* (Figure 1A, B, D). Importantly, these defects did not compromise cell viability in the γL262P bloodstream form, a mutant capable of surviving without kDNA, thereby affirming TAC53’s specific role in kDNA maintenance (Figure 1E).

Except for TAC53 several additional putative TAC candidates were identified in the SILAC RNAi experiments. However, none of them show the expected localization or RNAi phenotypes that would be expected for typical TAC subunits. This suggests that, within the limit of the proteomic analysis, all TAC subunits that function exclusively in the TAC have been discovered.

TAC53 localizes in close proximity to the kDNA, as observed via U-ExM, and occupies a position distal to TAC102 within the TAC hierarchy. Furthermore, the localization of TAC53 is TAC102-dependent, while TAC102 itself localizes independently of TAC53. Moreover, the abundance of HMG44 and KAP68 decreases upon depletion of TAC53 (Supplementary Figure S8 A, B) (35). Based on its position at the kDNA, TAC53, along with HMG44, KAP68, and KAP3, might mediate the interaction between the kDNA and the TAC. This model is further substantiated by the presence of canonical DNA-binding motifs within TAC53, including AT-hook and SPKK domains.

In replicating kDNA networks (1kdiv1N stage), TAC53 is conspicuously absent from the central region of the duplicated kDNA disc, an area predominantly composed of maxicircles that replicate immediately prior to segregation (54). This observation suggests that the interaction between TAC53 – and by extension the TAC – and the kDNA may be preferentially mediated through minicircles rather than maxicircles (Supplementary Figure S6C, D).

Although TAC53 shares functional parallels with other TAC proteins, it also displays distinct features. Notably, the relative abundance of TAC53 increases during mitochondrial genome replication, possibly reflecting a role in the reattachment of newly synthesized minicircles to the segregation apparatus. We propose a two-step model in which TAC53 and its interacting partners - HMG44, KAP68, and KAP3 - initially associate with replicated minicircles independently of upstream TAC components. In a second step, a subset of these TAC53-containing complexes that are bound to minicircles become integrated into the maturing TAC, while the remainder are targeted for degradation. This model is further supported by experimental evidence showing that early-stage TAC53 assembly is dependent on the presence of kDNA (Figure 4A, B), whereas this dependence diminishes significantly once TAC53 is integrated into the TAC (Figure 4C).

Collectively, we present a refined three-dimensional model of the tripartite attachment complex including its most distal component TAC53 and its interactors that mediate minicircle-specific tethering prior to integration into the mature TAC structure. The data strongly supports a model in which the mitochondrial genome is present in a diploid state and where each haploid genome is delineated by one TAC structure.

## Materials and Methods

### Cell lines

All procyclic cell (PCF) lines are derivates of the *T. brucei* single marker inducible 427 cell line described in (24). PCF cells were grown at 27°C in SDM-79 (55) and supplemented with 10% (v/v) fetal calf serum (FCS). Bloodstream form (BSF) cell lines are based on the New York single marker (NYsm) strain or on a derivative thereof named γL262P (36). BSF were grown in HMI-9 (56) containing 10% (v/v) FCS at 37 °C with 5% CO2. The tetracycline-inducible RNAi of p197 and TAC42 have already been described in (24) and (28), respectively. For tetracycline-inducible RNAi of the candidate genes, cells were stably transfected with a NotI-linearized pLew100-derivate plasmid containing stem-loop sequences covering parts of the ORF or the 3’UTR of the respective genes (see Supplementary Table S6 for specifications). C-terminally *in situ* HA- or myc tagging of the candidate genes was done by stable transfection of PCR products using plasmids of the pMOtag series (57) as templates with primers defining the sites of homologous recombination leading to expression of HA- or myc tagged full-length proteins.

### TAC depletomic assay using SILAC based MS analysis of isolated flagella

Cells capable of inducible RNAi against p197 or TAC42 were grown in SDM80 (58) supplemented with either light (^12^C6/^14^Nχ) or heavy (^13^C6/^15^Nχ) isotopes of arginine (1.1 mM) and lysine (0.4 mM) (Eurisotop) and 10% dialyzed FCS (BioConcept, Switzerland). To ensure complete labelling, cells were growing in this medium for 5 days in total. 3 days before harvesting, the cells growing in either light or heavy isotopes were induced with tetracycline for RNAi. Induced and uninduced cells were mixed in a 1:1 ratio, and 1x10^8^ cells in total were subtracted to flagellar preparation as described in (59). In brief, cells were harvested by centrifugation after EDTA (100nM) was added to the culture, directly resuspended in extraction buffer (10mM NaH2PO4, 150mM NaCl, 1mM MgCl2, pH 7.2) containing 0.5% Triton X-100 and incubated for 10 min on ice. After centrifugation pellets were incubated on ice for 45min with extraction buffer containing 1mM CaCl2. Extracted flagella could then be pelleted for 10min at 10’000g and were flash frozen in liquid nitrogen until used for liquid chromatography (LC)-MS analysis Experiments were performed in three (TAC42 RNAi) or four (p197 RNAi) biological replicates including a light/heavy label-switch. For LC-MS analysis, samples were thawed on ice and the extracted flagella were resuspended in urea buffer (8 M urea dissolved in 50 mM NH4HCO3). Cysteine residues were reduced by incubation with 5 mM Tris(2-carboxy-ethyl)phosphine/10 mM NH4HCO3 (30 min at room temperature) and free thiol groups were subsequently alkylated using 50 mM iodoacetamide/10 mM NH4HCO3 (30 min at room temperature in the dark). The reaction was quenched by addition of dithiothreitol at a final concentration of 33 mM. For tryptic in solution digestion, samples were diluted to approximately 1.6 M urea using 50 mM NH4HCO3 and proteins were digested overnight at 37°C. Peptide mixtures were desalted on StageTips (60), dried *in vacuo*, resuspended in 0.1% trifluoroacetic acid and analyzed on a Q Exactive (p197 SILAC RNAi samples) or an Orbitrap Elite (TAC42 SILAC RNAi samples) mass spectrometer (Thermo Fisher Scientific, Bremen, Germany), each coupled to an UltiMate 3000 RSCLnano HPLC system (Thermo Fisher Scientific, Dreieich, Germany). Proteins were identified and quantified using MaxQuant/Andromda (version 1.5.5.1; (61). MS/MS data were searched against all protein sequences for *T. brucei* TREU927 provided by the TriTrypDB database (https://tritrypdb.org; version 8.1) (47) using MaxQuant default settings except that only one unique peptide was required for protein identification. Lys8 and Arg10 were set as heavy labels and the options “requantify” and “match between runs” were enabled. Relative protein quantification was based on unique peptides and a minimum of one SILAC peptide pair. The mean log10 of normalized proteins abundance ratios (+/- Tet) was calculated, and for all proteins quantified in at least two replicates, p-values were determined using a two-sided Student’s t-test.

### Immunofluorescence Microscopy

For whole cell Immunofluorescence analysis cells were settled on a glass slide (Superfrost^TM^ Microscope Slides, Epredia) for 10 minutes, fixed with 4 % paraformaldehyde (PFA) for 10 minutes and permeabilized with 0.2 % Triton X-100 in phosphate-buffered saline (PBS) for 5 minutes. Cells were washed twice with PBS in between and blocked with 4 % bovine serum albumin (BSA) in PBS. Primary and secondary antibodies (see below for specifications) were diluted in PBS containing 2 % BSA and were incubated for 45 minutes in between PBS washes. After the last wash with PBS, DAPI prolong (Thermofisher, P36931) was added and sealed with a cover glass.

For flagellar extraction, the cells were harvested by centrifugation after adding EDTA to a final concentration of 5 mM to the culture. The cells were pelleted and resuspended in extraction buffer (10 mM NaH₂PO₄, 150 mM NaCl, 1 mM MgCl₂, pH 7.2) containing 0.5% Triton X-100 and incubated on ice for 10 minutes. After centrifugation, the pellets were incubated on ice for 45 minutes in extraction buffer supplemented with 1 mM CaCl₂. The extracted flagella were pelleted by centrifugation at 3000 × g for 5 minutes at 4°C, resuspended in PBS, settled onto a coverslip for 10 minutes, and fixed with 4% PFA. After fixing, the described Immunofluorescence protocol was applied.

Immunofluorescence assay images for whole cells and extracted flagella were acquired on a LEICA DM5500B Widefield Microscope with a 100x (1.4 NA) objective and analyzed in ImageJ.

### TAC53 sequence analysis

Protein sequences and accession numbers for TAC used in this study were retrieved from the TriTryp database (47). Searches for TAC53 homologs were done using BLAST in the TriTryp database or using hmmsearch using custom hmm profiles (HMMER version 3.0) (47, 62). Multiple sequence alignment was performed with MAFFT (L-INS-i method, version 7) and visualized with the Clustalx coloring scheme in Jalview (version 2.11) (63, 64). AT hooks and SPKK motifs were identified by manual inspection (51, 52).

### U-ExM

Coverslips (12 mm, CB00120RA120MNZ0, Epredia) were functionalized with poly-D-lysine (A3890401, Gibco) at room temperature for 30 minutes, followed by three washes with deionized water. Stained cells (prepared as per the immunofluorescence assay protocol) were spread onto the functionalized coverslips and allowed to settle at room temperature in the dark for 10 minutes.

Cells were anchored using 0.7% formaldehyde (FA, 36.5–38%, F8775, SIGMA) and 1% acrylamide (AA, 40%, A4058, SIGMA) in PBS at 37°C for 2 hours. Then, gelation was performed with a freshly prepared gelation solution consisted of 19% sodium acrylate (SA, 97–99%, 408220, SIGMA), 10% acrylamide (AA, 40%, A4058, SIGMA), and 0.1% N,N’-methylenebisacrylamide (BIS, 2%, M1533, SIGMA), supplemented with 0.5% ammonium persulfate (APS, 17874, Thermo Fisher) and 0.5% tetramethylethylenediamine (TEMED, 17919, Thermofisher). Gelation was carried out on ice. A 35 µl drop of gelation solution was placed on parafilm in a precooled humidity chamber, and cells were incubated on ice for 5 minutes, followed by final gelation at 37°C for 30 minutes.

Gels were detached from the coverslips by gentle shaking in 1 ml of denaturation buffer (200 mM SDS, 200 mM NaCl, and 50 mM Tris in deionized water, pH 9) at room temperature for 15 minutes. Gels were then incubated in the denaturation buffer at 95°C for 30 minutes. Afterward, gels were washed three times for 5 minutes in PBS, and DNA was stained with 5 µg/ml DAPI (D9542-5MG, SIGMA) in PBS (diluted 1:1000) with gentle agitation for one hour. Samples were then expanded in deionized water overnight.

### Mounting and Acquisition for ExM images

The gel was cut and mounted on poly-D-lysine functionalized 25 mm glass-bottom dishes (D35-20-1.5-N, Cellvis, 35 mm glass bottom dish with 20 mm microwell #1.5 cover glass). Z-stacks were acquired using a Nikon Ti2 Kinetix Widefield Microscope (equipped with a Photometrics Kinetix 500 fps camera) on a 100x (NA =1.45) objective. Following parameters were used: z step size was set to 0.2 µm and pixel size was 65 nm. Images were deconvolved using Huygens software HRM and analyzed with ImageJ. To quantify the images, z – projections were done using ImageJ with Maximum Intensity as Projection type. Measurements of TAC diameters were analyzed using GraphPad Prism version 9.5.0 (www.graphpad.com). Statistical analysis was performed using an Ordinary One-way ANOVA using the Bonferroni multiple comparisons test (ns = non-significant, *P ≤ 0.05; **P ≤ 0.01; ***P ≤ 0.001; ****P ≤ 0.0001).

### Immunoprecipitation

Our results indicate that sonication is the most effective method for solubilizing kDNA-proximal proteins, outperforming chemical fractionation methods like digitonin or RIPA buffer, as shown for KAP68 and HMG44 (Supplementary Figure S9). Mitochondria-enriched pellets (with 0.015% digitonin in SoTe (20 mM Tris HCl pH 7.5, 0.6 M sorbitol, 2 mM EDTA)) (5 x 10^7^ cells each) were sonicated using a Bioruptor Plus sonication device by diagenode. Sonication was performed on level H for 10 times 30 seconds on/off at 4°C. Samples were then centrifuged at 4 ° C for 20 minutes and supernatants were applied to either anti-c myc (ETview Red Anti-myc Affinity Gel, Sigma-Aldrich, Merck, E6654) or anti-HA beads (EZview Red Anti-HA Affinity Gel, Sigma-Aldrich, Merck, E6779) in an overnight incubation at 4° C. Beads were then washed 3 times in bead wash buffer (2x lysis buffer (40mM Tris-HCL, ph 7.4, 0.2mM EDTA, 200mM NaCl, 50mM KCl, H2O), 25x cOmplete, Mini, EDTA-free Protease Inhibitor Cocktail (Sigma-Aldrich, Merck, 04693159001) and H2O, centrifuged and sent for MS analysis.

### Antibodies

For this study, the following primary and secondary antibodies were used (1:1000 for Immunofluorescence assays; 1:500 for Expansion Microscopy): Anti-HA.11 Epitope Tag Antibody, mouse (BioLegend, MMS-101R); Anti-c-Myc Monoclonal Antibody, mouse ((Thermofisher, 13-2500); Anti-c-Myc antibody, rabbit ((SigmaAldrich, C3956); HA tag antibody, rabbit (Thermofisher, 71-5500); Goat anti-Mouse IgG (H+L) Cross-Adsorbed Secondary Antibody, Alexa Fluor™ 488 (Thermofisher, A-11001); Goat anti-Mouse IgG (H+L) Cross-Adsorbed Secondary Antibody, Alexa Fluor™ 594 (Thermofisher, A-11005); Goat anti-Rabbit IgG (H+L) Cross-Adsorbed Secondary Antibody, Alexa Fluor™ 488 (Thermofisher, A-11008); Goat anti-Rabbit IgG (H+L) Cross-Adsorbed Secondary Antibody, Alexa Fluor™ 594 (Thermofisher, A-11012).

The TAC53 antibody was produced at Eurogentec targeting the following peptide sequence: 473-487 (H - CAT TPQ RIT NTG LLR L – OH).

### Northern blots

Total RNA was extracted using acid guanidinium thiocyanate-phenol-chloroform according to (65) and separated on a 1% agarose gel. Northern blot probes for the corresponding RNAi targeting fragment were radioactively labelled using the Prime-a-Gene labelling system (Promega). Immunoblots- and Northern blots were done as previously described (66).

### MS sample preparation for immunoprecipitation assays

The samples were processed via in-gel digestion as described previously (67), whereby denatured proteins were run on a 4–12% NuPage Bis-/Tris Gel (Thermo Scientific) at 180 volts for 8 min in 1x MES SDS Running Buffer (Thermo Scientific). Proteins in the gel were fixed with MS fixation buffer (7% acetic acid, 40% methanol) and stained with Coomassie Blue (10% acetic acid, 45% ethanol, 0.25% w/v brilliant blue (Roth)) for 5 minutes. The gels were destained overnight in water. The next day, each sample of the gel was cut into 1x1 mm squares, and the gel pieces were further destained with 50% ethanol and 25 mM f.c. ammonium bicarbonate digestion buffer (ABC pH 8.0) for several rounds and dehydrated with acetonitrile. Disulfide bridges were reduced with 10 mM f.c. DTT in 50 mM ABC at 56°C for 60 min and alkylated with 50 mM f.c. iodoacetamide in 50 mM ABC at 25°C for 45 min in the dark. The gel pieces were washed with 50 mM ABC and dehydrated three times with acetonitrile. The proteins in the gel pieces were subsequently digested overnight with 1 µg of MS-grade trypsin (Serva) in 50 mM ABC (pH 8.0) at 37°C. The next day, the digested proteins were extracted from the gel pieces with two rounds of extraction buffer (30% acetonitrile in 50 mM ABC buffer) and dehydrated with three rounds of acetonitrile for 15 min at 25°C. The supernatants were combined, and the samples were placed in a concentrator to evaporate the acetonitrile until the volume was <200 µl. They were then desalted by StageTipping as described previously (60), where two layers of Empore C18 material (3M) were pushed into a 200 µl pipette tip, activated with 50 µl of methanol, equilibrated with 50 µl of Buffer B (80% acetonitrile, 0.1% formic acid), and washed with 50 µl of Buffer A (0.1% formic acid). The samples were added and washed with Buffer A. To run on the mass spectrometer, samples were eluted from the stage tip with 30 µl of Buffer B, placed in a concentrator to remove the acetonitrile until the remaining volume was 7 µl, and diluted with Buffer A for a final volume of 14 µl.

### MS measurement and data analysis for immunoprecipitation assays

Five microliters of sample were injected on an Easy nLC-1000 (Thermo Scientific) coupled to a Q Exactive Plus (Thermo Scientific). On the Easy nLC-1000, tryptic peptides were separated through reversed-phase liquid chromatography on a 20 cm column with 75 µm inner diameter, (New Objective) self-packed with ReproSil-Pur 120 C18-AQ 1.9 (Dr. Maisch GmbH), mounted on an electrospray ion source. The samples were eluted from the column at a 225 nL*min^-1^ flow rate with a 105 min gradient ranging from 2–95% Buffer B (0.1% formic acid, 80% acetonitrile). Peptides were sprayed into the Q Exactive Plus set to positive ion mode operated with a TOP10 data-dependent acquisition scheme. The full scan was conducted with a resolution of 70,000, and the MS/MS scans had a resolution of 17,500. The raw MS data were processed with MaxQuant version 2.4.2.0 with the default settings. Proteins were quantified via the LFQ quantification algorithm (without FastLFQ) with 2 LFQ ratio counts unique + razor peptides. The false discovery rate was set to 0.01. Contaminants, protein groups identified only by site, reverse database hits, and proteins with fewer than two peptides removed prior to further analysis. Missing values were imputed.

## Data availability

The mass spectrometry proteomics data have been deposited to the ProteomeXchange Consortium (68) via the PRIDE (69) partner repository and are accessible using the dataset identifiers PXD063227 (p197 SILAC RNAi), PXD063233 (TAC42 SILAC RNAi), PXD061646 (KAP68-IP) and PXD062506 (TAC53- and HMG44-IPs).

## Acknowledgments

Research in the lab of T. O. was supported by the Swiss National Science Foundation grants (179454 and 207525) and the Uniscientia foundation. Work in the lab of A.S. was supported in part by NCCR RNA & Disease, a National Centre of Competence in Research (grant number 205601) and by project grant SNF 205200 both funded by the Swiss National Science Foundation. Work in the lab of B.A. was supported by a Wellcome Discovery Award (227243/Z/23/Z).

We want to thank the Microscopy Imaging Center (MIC) of the University of Bern and the team of the Proteomics and Mass Spectrometry Core Facility (PMSCF) at the Department for BioMedical Research (DBMR) for their valuable service. We additionally want to thank Bianca Berger and Markus Gerber for fruitful discussions regarding the topic.

