## Supporting Information for "Beyond a Linear Structure: The Tubular Organization of the Tripartite Attachment Complex and the Functional Role of TAC53"

**This PDF file includes:**

Figures S1 to S9
Tables S1 to S6

**A**

| gene | fold change p197 RNAi | annotation | kDa / pI | additional info | Tryptag | method tested |
|---|---|---|---|---|---|---|
| Tb927.10.13600 | 8.5 | CLDP18 | 18kDa / 11.7 | predicted TMD, IM cluster | mitochondrion | RNAi in γL262P |
| Tb927.5.2970 | 7.1 | TSIF | 90kDa / 4.5 | | mBB / pBB | RNAi in γL262P |
| Tb927.5.2560 | 2.4 | protein of unknown function | 9kDa / 6.5 | predicted TMD, IM cluster | cytoplasm | RNAi in γL262P |
| Tb927.10.6200 | 2.3 | protein of unknown function | 12kDa / 10.4 | predicted TMD | mitochondrion | RNAi in γL262P |
| Tb927.11.1700 | 2.2 | protein of unknown function | 62kDa / 4.0 | | kDNA | RNAi in PCF |
| Tb927.7.7090 | 2 | protein of unknown function | 17kDa / 11.6 | predicted TMD | mitochondrion | RNAi in γL262P |
| Tb927.5.2630 | 1.8 | protein of unknown function | 27kDa / 4.7 | predicted TMD | mitochondrion | RNAi in γL262P |

**B**

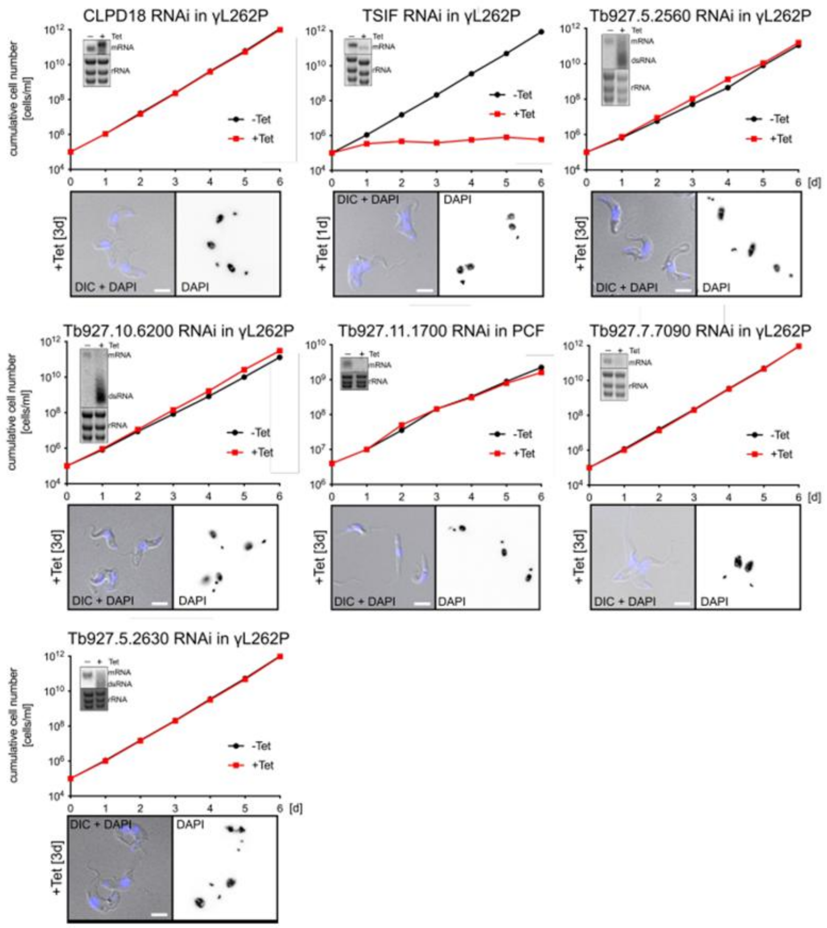


**Fig. S1. Analysis of Candidates from the p197-Depletome Assay**

A) Table of candidate proteins depleted by more than 1.5-fold upon p197-RNAi induction. The table includes accession number, fold change, annotation, molecular weight (kDa), isoelectric point (pI), predicted domains, their localization according to TrypTag.org, and the tested method. B) Growth curves (GC, upper panel) over 6 days for uninduced (-Tet, black) and RNAi-induced (+Tet, red) cells, along with DAPI stainings (lower panel) of the corresponding tested candidates. The tested gene and cell type are indicated above the GC. Time points of DAPI stainings are marked on the left of the images. GC inset: RNAi efficiency was tested using a northern blot (NB) probed against the respective RNAi target. Ethidium bromide-stained rRNAs serve as the loading control. Scale bar is 5 μm.

**A**

| gene | fold change TAC42 RNAi | annotation | kDa / pI | additional info | Tryptag | method tested |
|---|---|---|---|---|---|---|
| Tb927.10.8980 | 2.1 | protein of unknown function | 18kDa / 12.3 | IM cluster, only trypanosomal | kDNA | IFA, RNAi in PCF |
| Tb927.1.2730 | 1.8 | protein of unknown function | 39kDa / 6.6 | SET-domain | mitochondrion / kDNA | IFA, RNAi in PCF |
| Tb927.8.3160 | 1.7 | protein of unknown function | 17kDa / 10.4 | | cytoplasm / kDNA | IFA, RNAi in PCF |
| Tb927.10.11060 | 1.5 | protein of unknown function | 23kDa / 9.2 | | mitochondrion | IFA |
| Tb927.9.3340 | 1.5 | CMC1-like | 20kDa / 6.8 | IM cluster | cytoplasm | IFA |
| Tb927.7.6660 | 1.5 | chaperone protein DNAj, putative | 30kDa / 9.2 | DNAj domain | mitochondrion | IFA |
| Tb927.10.4240 | 1.5 | protein of unknown function | 15kDa / 10.1 | IM cluster | cytoplasm | IFA |

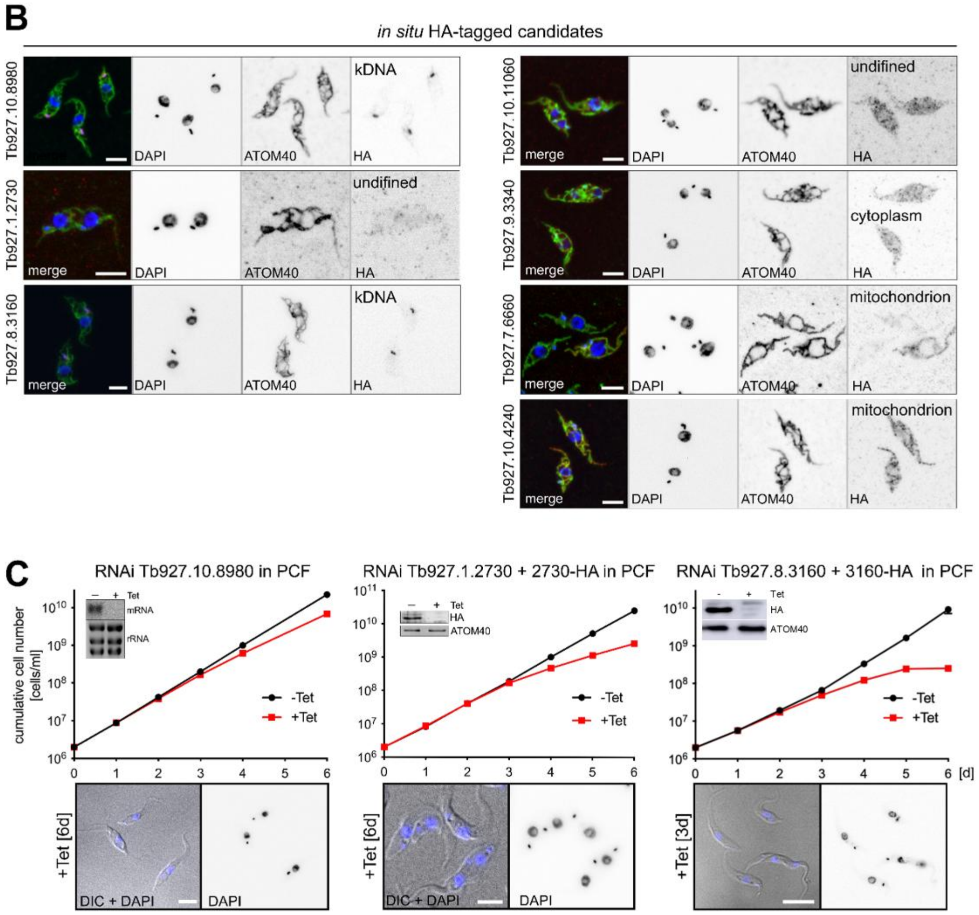


**Fig. S2. Analysis of Candidates from the TAC42-Depletome Assay**

A) Table of candidate proteins depleted by more than 1.5-fold upon TAC42-RNAi induction. The table includes accession number, fold change, annotation, molecular weight (kDa), isoelectric point (pI), predicted domains, their localization according to TrypTag.org, and the tested method.
B) Immunofluorescent images showing the localization of the respective candidate genes. Cells were stained with DAPI (blue), anti-ATOM40 (green) as a mitochondrial marker, and anti-HA (red) for the in-situ HA-tagged candidate proteins. Localization patterns are indicated in the images showing the HA staining. Scale bar: 5 µm.
C) Growth curves (upper panel) over 6 days of uninduced (-Tet, black) and RNAi-induced (+Tet, red) cells, along with DAPI stainings (lower panel) of the corresponding tested candidates. The tested gene and cell type are indicated above the growth curve. Time points of DAPI stainings are

marked on the left of the images. Growth curve left inset: RNAi efficiency was tested using a northern blot (NB) probed against the respective RNAi target. Ethidium bromide-stained rRNAs serve as the loading control. Growth curve middle and right inset: RNAi efficiency has been tested using a WB probed against the RNAi target and ATOM40 as loading control. Scale bar is 5 µm.

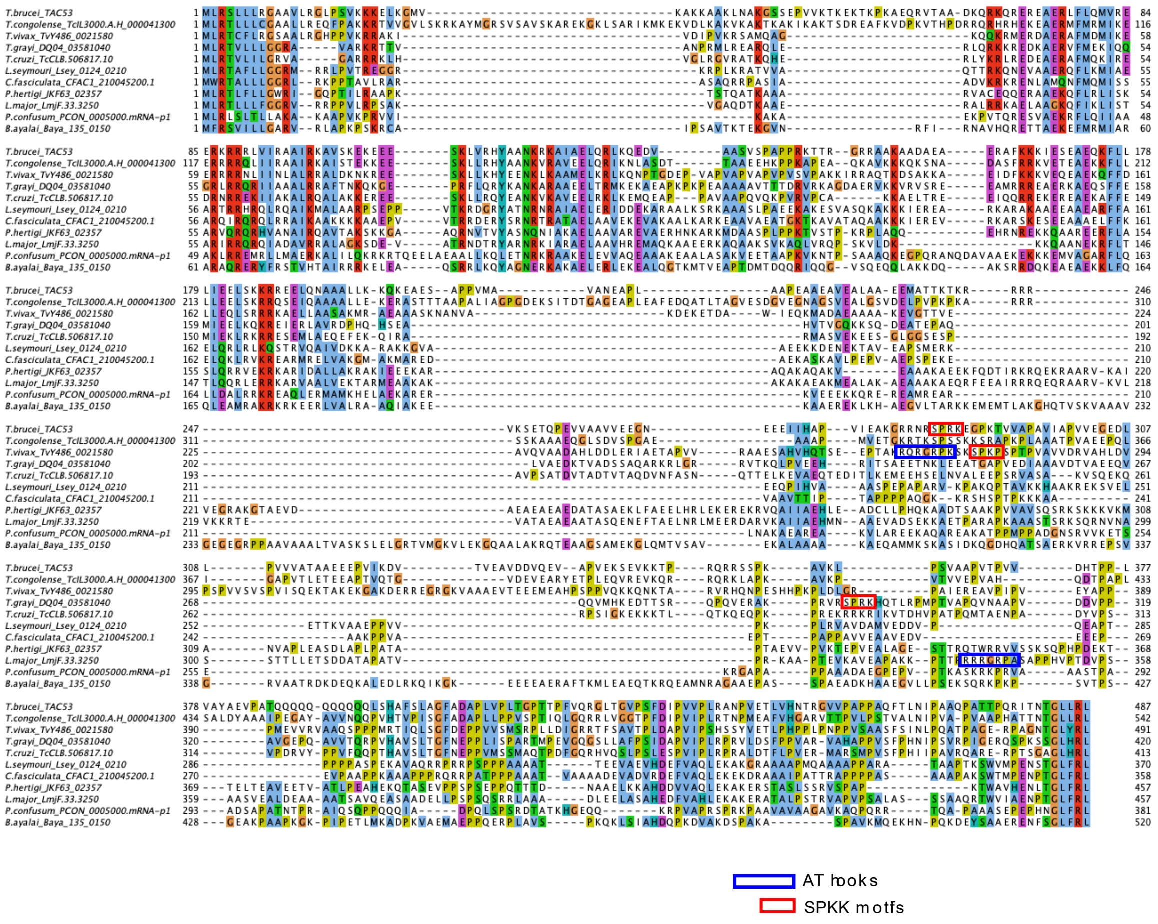


**Fig. S3. Multiple Sequence Alignment of TAC53 Homologs in Trypanosomatids**

The alignment highlights the presence of AT-hooks and SPKK motifs in some TAC53 homologs. AT-hooks are shown in blue, while SPKK motifs are shown in red.

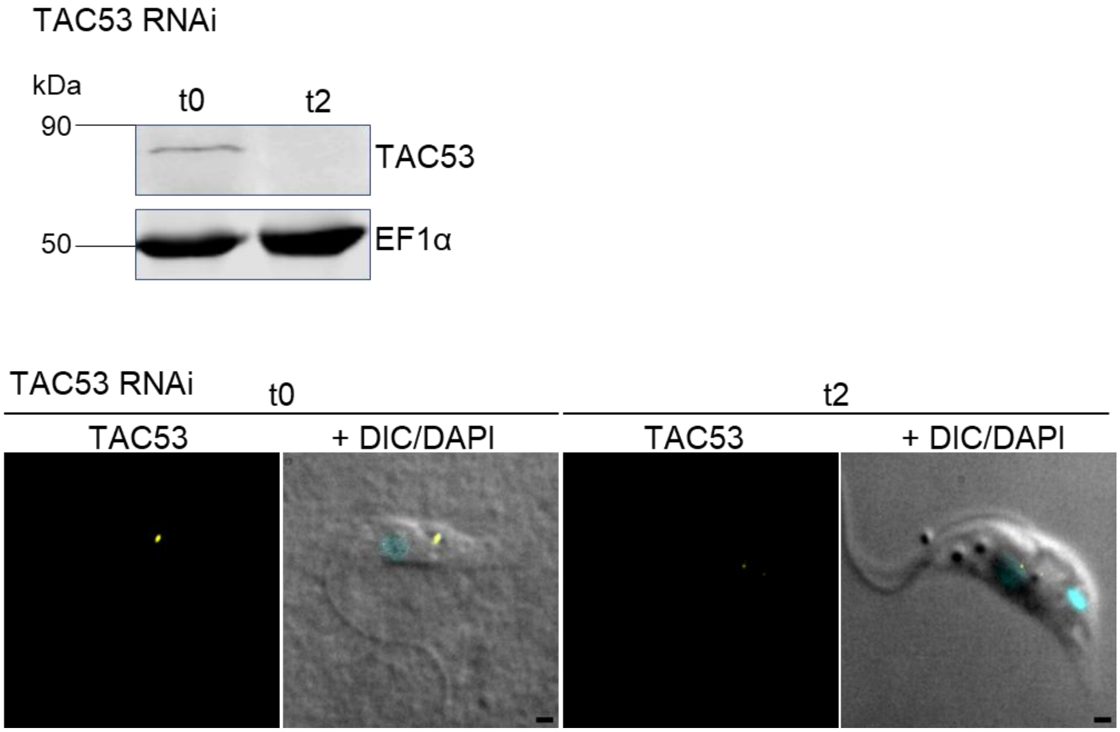


**Fig. S4. The TAC53 Antibody Specifically Binds TAC53 in Western Blot and Immunofluorescence Assays**

A) Western blot showing TAC53 signal in RNAi-uninduced and induced cells [2d]. EF1α is used as a loading control.

B) Immunofluorescence assay showing TAC53 signal in RNAi-uninduced and induced cells [2d]. TAC53 antibody signal is shown in yellow, and DAPI staining is shown in cyan. Scale bar is 1 μm.

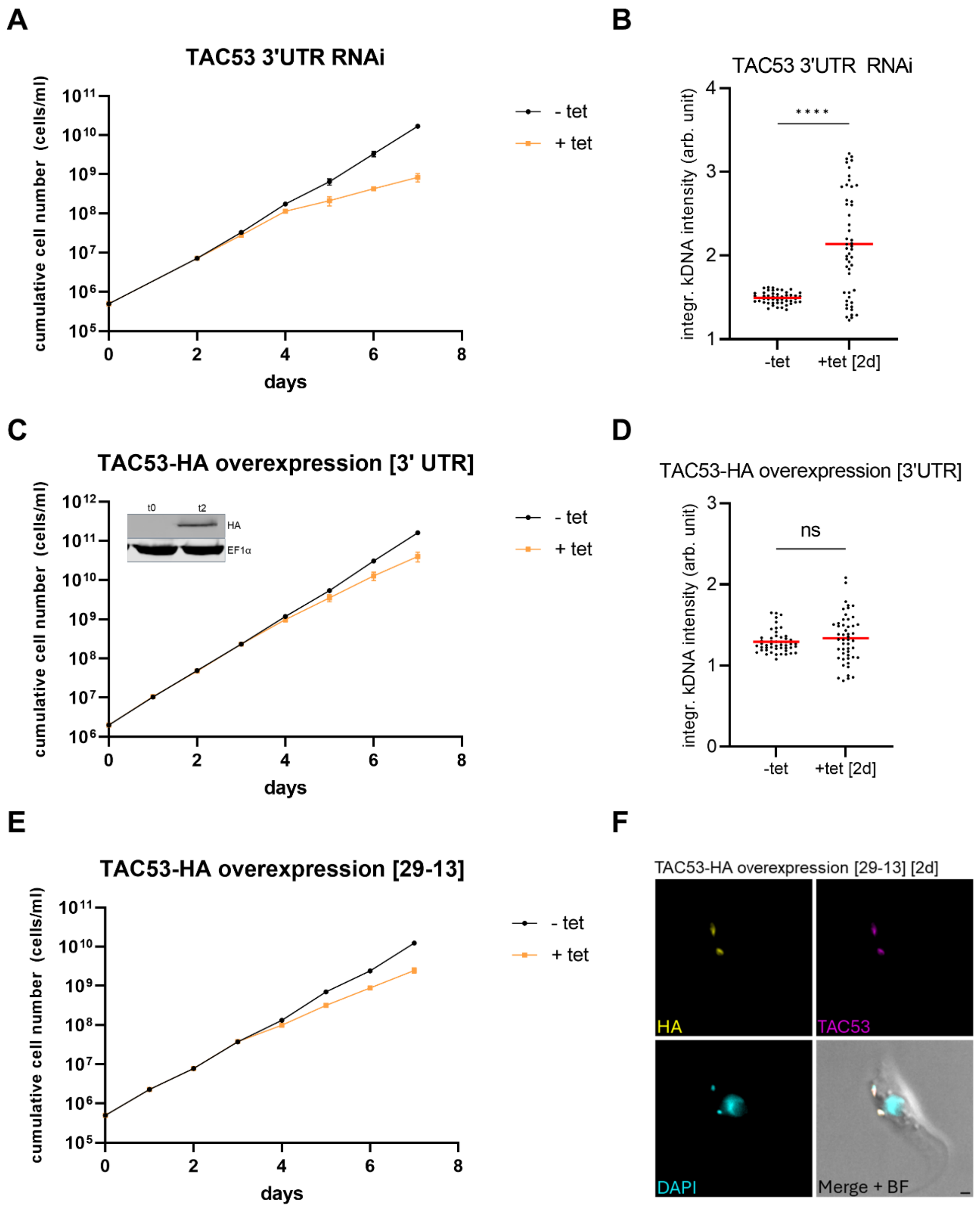


**Fig. S5. Analysis of TAC53 RNAi and Overexpression**

A) Growth curve of procyclic form cells targeting the 3' UTR of TAC53. Black line: uninduced cells; orange line: induced cells.

B) Integrated kDNA intensities measured in uninduced and induced [2d] cells from panel A. The red line marks the mean.

C) Growth curve of TAC53-HA expression in the background of the 3' UTR RNAi (exclusive

expression). Black line: uninduced cells; orange line: induced cells. Inset: Western blot showing HA signal after two days of induction, with EF1α as the loading control.
D) Integrated kDNA intensities measured in uninduced and induced [2d] cells from panel C.
E) Growth curve of TAC53-HA overexpression in 29-13 cells. Black line: uninduced cells; orange line: induced cells.
F) Immunofluorescence analysis of TAC53-HA overexpression. The HA tag is shown in yellow, the TAC53 antibody signal is shown in magenta, and DAPI is shown in cyan. BF=Brightfield. Scale bar is 1 μm.

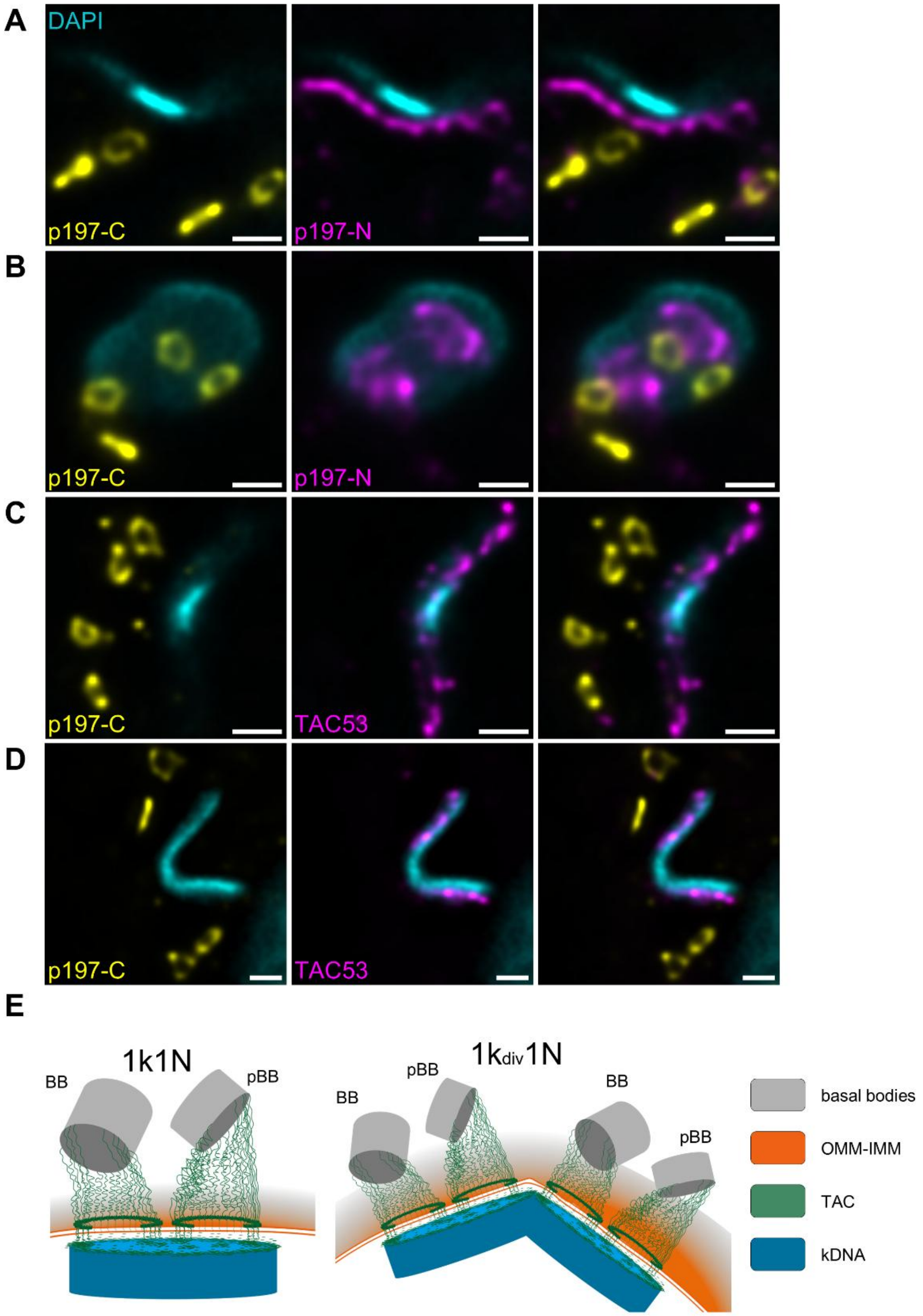


**Fig. S6. Expansion Microscopy Images of Replicated kDNAs Displaying Four TAC Structures**

A) Lateral perspective of replicated kDNAs showing the p197 C- and N-terminus. The C-terminus is shown in yellow, the N-terminus in magenta, and DAPI is shown in cyan. The scale bar is 1µm (not adjusted for Expansion).

B) Replicated kDNA shown from an axial perspective with p197 C-terminus in yellow and p197 N-terminus in magenta. DAPI is shown in cyan.
C) D) Replicated kDNA shown from two different lateral perspectives with p197 C-terminus in yellow, TAC53 in magenta, and DAPI in cyan. Scale bar for A-D is 1 µm.
(E) Illustration of the observed structure in expansion microscopy showing pro- and basal bodies (pBB and BB), outer- and inner mitochondrial membranes (OMM-IMM), the TAC, and the kDNA in 1k1N and 1kdiv1N cells. Color coding is found in the figure.

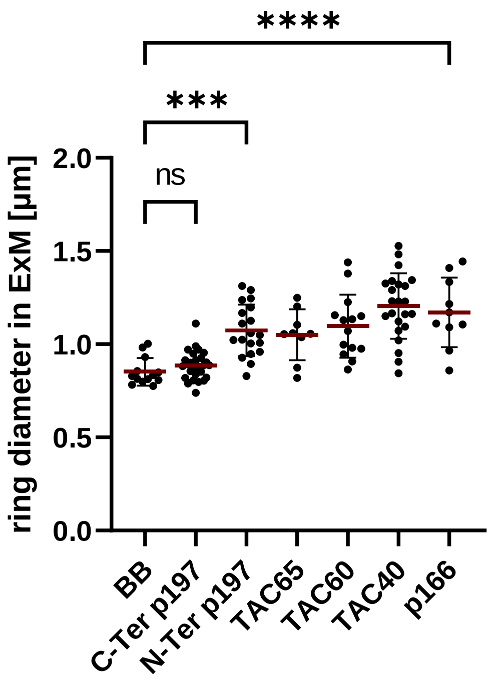


**Fig. S7. Ring diameter measurements of the basal body and TAC components in Expansion Microscopy.**

Diameters of ring-like structures were measured from Expansion Microscopy images for the basal body and several TAC proteins, including the C- and N-terminus of p197, TAC65, TAC60, TAC40, and p166. The y-axis shows the measured ring diameters in micrometers (µm), and the x-axis indicates the specific proteins. Each data point represents an individual measurement. Red line marks the mean. Statistical analysis was performed using GraphPad Prism with ordinary one-way ANOVA followed by Bonferroni's multiple comparison test.

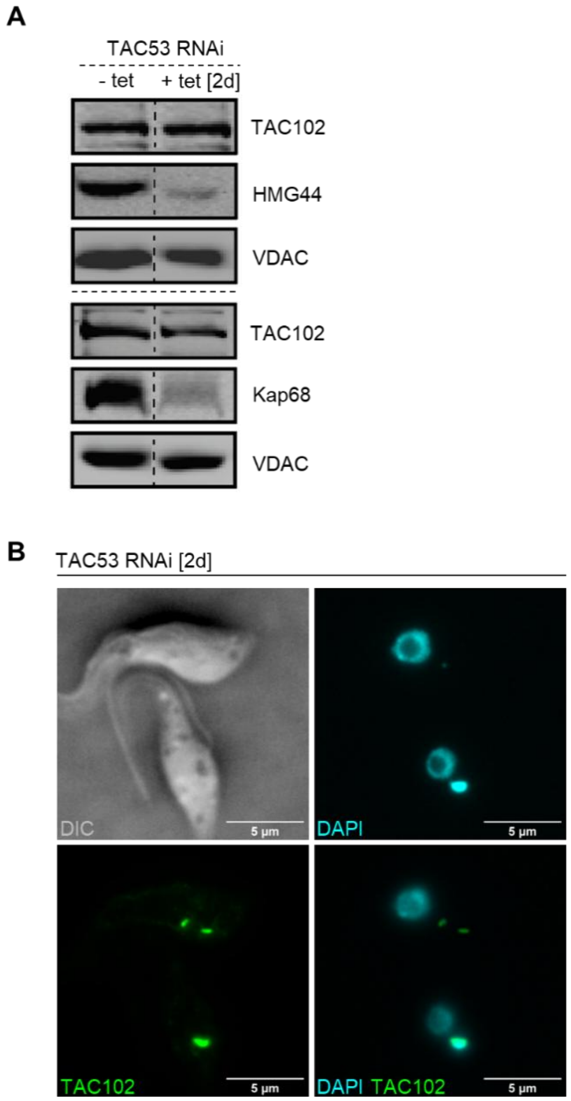


**Fig. S8. Protein dependencies.**

A) HMG44 and KAP68 are depleted after TAC53 RNAi [2d], while TAC102 remain unchanged. Western Blot analysis of TAC102, HMG44 and KAP68 signals following TAC53 depletion. VDAC serves as a loading control.

B) Immunofluorescence analysis of TAC53-depleted cells [2d], showing TAC102 signals in green, DAPI in cyan, and DIC. Scale bar is 5 µm.

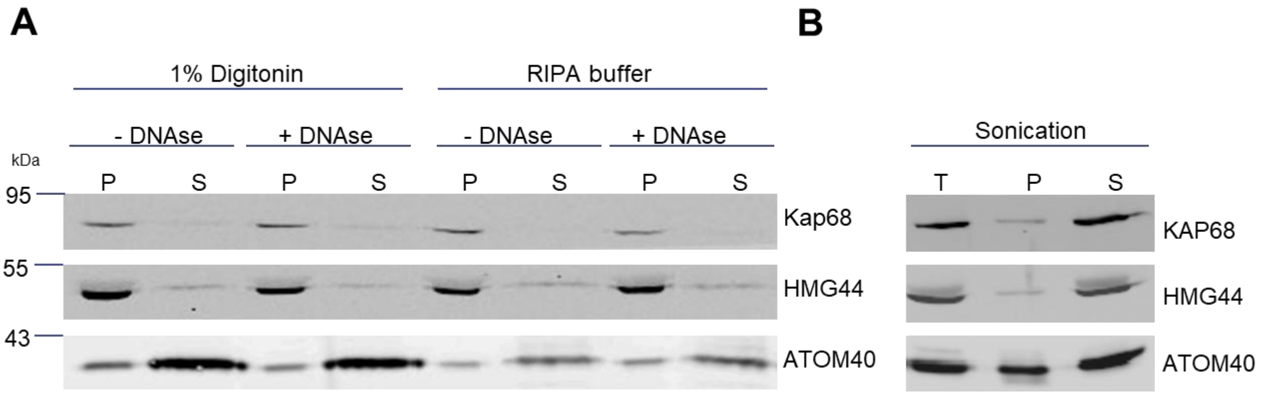


**Fig. S9. Solubilization Assay Comparing Sonication and Chemical Fractionation for HMG44 and KAP68**

A) Western blot analysis showing pellet and supernatant fractions for HMG44 and KAP68 after treatment with 1% digitonin or RIPA buffer. Enriched mitochondria were either treated with DNase-I or left untreated.
B) Western blot analysis showing total cells, pellet, and supernatant after sonication. ATOM40 serves as the loading control.

**Table S1. p197-Depleteome List**

All proteins downregulated by more than 1.5-fold upon p197 depletion are listed. The table includes accession number, gene name, depletion factor, statistical significance (p-value ≤ 0.05), TritrypDB annotation, localization according to TrypTag.org, and indication of detection in the TAC42-RNAi dataset. Color coding as follows: TAC proteins (red), BB proteins (blue), common candidates (green), p197-exclusive candidates (orange), and COX proteins (gray).

| acc. number | gene name | fold change (-Tet/+Tet) | p-value ≤ 0.05 | protein function / feauters / Tryptag localization | found in TAC42 RNAi data set |
|---|---|---|---|---|---|
| Tb927.10.15750 | p197 | 11.9 | ✓ | TAC | ✓ |
| Tb927.10.13600 | CLDP18 | 8.5 | | protein of unknown function, IM cluster | ✓ |
| Tb927.5.2970 | TSIF | 7.1 | | suppressive immunomodulating factor, mBB/pBB localization | |
| Tb927.11.12040 | | 3.7 | | protein of unknown function, putative cytochrome C, mitochondrion | ✓ |
| Tb927.2.6100 | | 3.6 | ✓ | protein of unknown function, kDNA | ✓ |
| Tb927.11.3290 | p166 | 3.5 | ✓ | TAC | ✓ |
| Tb927.11.16580 | | 3.3 | | protein of unknown function, TPR-like domain, FP | ✓ |
| Tb927.5.830 | TAC65 | 3.1 | ✓ | TAC | ✓ |
| Tb927.9.5020 | mtHMG44 | 2.7 | ✓ | protein of unknown function, kDNA | ✓ |
| Tb11.0400 | P27 | 2.5 | ✓ | P27 protein | ✓ |
| Tb927.5.2560 | | 2.4 | ✓ | protein of unknown function, IM cluster, TMD predicted | ✓ |
| Tb927.4.4620 | COXVIII | 2.4 | ✓ | cytochrome oxidase subunit VIII (COXVIII) | ✓ |
| Tb927.10.14860 | NDUFA6 | 2.4 | | NADH-ubiquinone oxidoreductase B14 subunit, mitochondrion | ✓ |
| Tb927.10.6200 | | 2.3 | ✓ | protein of unknown function, TMD predicted | ✓ |
| Tb927.10.280 | COXVI | 2.3 | ✓ | cytochrome oxidase subunit VI (COXVI) | ✓ |
| Tb927.10.8320 | COXIX | 2.2 | ✓ | cytochrome oxidase subunit IX (COXIX) | ✓ |
| Tb927.11.16120 | KAP68 | 2.2 | ✓ | protein of unknown function, kDNA associated | ✓ |
| Tb927.9.3170 | COXV | 2.2 | ✓ | cytochrome oxidase subunit V (COXV) | ✓ |
| Tb927.11.1700 | | 2.2 | ✓ | protein of unknown function | ✓ |
| Tb927.11.16400 | KAP3 | 2.1 | ✓ | kinetoplast DNA-associated protein | ✓ |
| Tb927.3.1410 | COXVII | 2.1 | | cytochrome oxidase subunit VII (COXVII) | ✓ |
| Tb927.7.7090 | | 2.0 | | protein of unknown function | ✓ |
| Tb927.11.11480 | | 2.0 | | Trichohyalin, FAZ | ✓ |
| Tb927.1.4100 | COXIV | 1.9 | ✓ | cytochrome oxidase subunit IV (COXIV) | ✓ |
| Tb927.11.1130 | | 1.9 | ✓ | calpain-like cysteine peptidase, putative, cytoplasm | ✓ |
| Tb927.7.2390 | TAC102 | 1.9 | ✓ | TAC | ✓ |
| Tb927.7.1400 | TAC60 | 1.9 | ✓ | TAC | ✓ |
| Tb927.5.2630 | | 1.8 | | protein of unknown function, TMD predicted | |
| Tb927.10.9080 | | 1.8 | ✓ | pteridine transporter, putative, cytoplasm and pellicular membrane | ✓ |
| Tb927.8.3630 | FT2 | 1.8 | ✓ | folate transporter, putative, cytoplasm | ✓ |
| Tb927.10.4280 | | 1.7 | ✓ | ubiquinol-cytochrome c reductase complex, mitochondrion | ✓ |
| Tb927.4.1610 | TAC40 | 1.7 | ✓ | TAC | ✓ |
| Tb927.11.8430 | | 1.7 | ✓ | protein of unknown function | ✓ |
| Tb927.5.2790 | Pol beta-PAK | 1.7 | ✓ | mitochondrial DNA polymerase beta-PAK (Pol beta-PAK), antipodal sites | ✓ |
| Tb927.10.13310 | RPB5z | 1.7 | | DNA-directed RNA polymerase I subunit, cytoplasm, and nucleus | ✓ |
| Tb927.6.2760 | BBP164 | 1.6 | | BB protein | ✓ |
| Tb927.7.5700 | pATOM36 | 1.6 | ✓ | TAC, mitochondrial import machinery | ✓ |
| Tb10.v4.0166 | | 1.6 | | variant surface glycoprotein (VSG, pseudogene) | |
| Tb927.7.3060 | TAC42 | 1.6 | ✓ | TAC | ✓ |
| Tb927.11.10370 | | 1.5 | ✓ | glycosyl hydrolase-like protein, golgi | ✓ |
| Tb927.8.3650 | FT3 | 1.5 | ✓ | folate transporter, ER | ✓ |
| Tb927.11.10160 | | 1.5 | | 60S ribosomal protein L22, cytoplasm | ✓ |
| Tb927.10.230 | | 1.5 | | proteasome alpha 5 subunit, cytoplasm and nucleus | |

**Table S2. TAC42-Depleteome List**

All proteins downregulated by more than 1.5-fold upon TAC42 depletion are listed. The table includes accession number, gene name, depletion factor, statistical significance (p-value ≤ 0.05), TritrypDB annotation, localization according to TrypTag.org, and indication of detection in the p197-RNAi dataset. Color coding as follows: TAC proteins (red), BB proteins (blue), common candidates (green), TAC42 exclusive candidates (pink).

| acc. number | gene name | fold change (-Tet/+Tet) | p-value ≤ 0.05 | protein function / feauters / Tryptag localization | found in p197 RNAi data set |
|---|---|---|---|---|---|
| Tb927.7.1400 | TAC60 | 14.4 | ✓ | TAC | ✓ |
| Tb927.7.3060 | TAC42 | 13.5 | | TAC | ✓ |
| Tb927.11.3290 | p166 | 8.3 | ✓ | TAC | ✓ |
| Tb927.11.11590 | EIF3E | 8.2 | | eukaryotic translation initiation factor, putative, cytoplasm | |
| Tb927.9.2630 | | 6.5 | | protein of unknown function | |
| Tb927.2.6100 | | 5.6 | ✓ | protein of unknown function | ✓ |
| Tb927.9.5020 | HMG44 | 4.4 | ✓ | protein of unknown function | ✓ |
| Tb927.10.13870 | | 3.8 | | tubulin tyrosine ligase protein, putative, cytoplasm and flagellum | |
| Tb927.6.200 | | 3.7 | | receptor-type adenylate cyclase GRESAG 4 | |
| Tb927.11.6870 | 14-3-3-II | 3.6 | | 14-3-3 protein, FP and flagellum and cytoplasm | |
| Tb927.7.7400 | BBP248 | 3.2 | | BB protein | ✓ |
| Tb927.8.2770 | IP3R | 2.6 | | insulin receptor, Ryanodine receptor, ER | |
| Tb927.11.180 | | 2.5 | | electron transfer flavoprotein, putative, mitochondrion and kDNA | ✓ |
| Tb927.7.2390 | TAC102 | 2.5 | ✓ | TAC | ✓ |
| Tb927.10.1280 | | 2.5 | | Domain of unknown function (DUF4499), putative, ER | |
| Tb927.11.3620 | NT8.2 | 2.4 | ✓ | nucleobase/nucleoside transporter 8.1 (NT8.1), ER | |
| Tb927.7.320 | RBP8 | 2.4 | | RNA-binding protein 8, cytoplasm | |
| Tb927.7.4440 | | 2.2 | ✓ | NAD dependent epimerase/dehydratase family, putative, mitochondrion and kDNA | |
| Tb927.10.8980 | | 2.1 | ✓ | protein of unknown function | |
| Tb927.9.8620 | | 2.1 | | protein of unknown function, cytoplasm | |
| Tb927.10.14400 | | 2.1 | | protein of unknown function, FAZ | |
| Tb927.5.830 | TAC65 | 2.0 | ✓ | TAC | ✓ |
| Tb927.5.3240 | | 2.0 | | protein of unknown function, nuclear protein | |
| Tb927.2.5870 | | 1.9 | | protein of unknown function, cytoplasm and flagellar tip | ✓ |
| Tb927.7.4970 | GS | 1.8 | | glutamine synthetase, putative | |
| Tb927.1.2730 | | 1.8 | ✓ | protein of unknown function | |
| Tb927.10.9810 | | 1.8 | ✓ | protein of unknown function | |
| Tb927.10.5920 | | 1.7 | | protein of unknown function, nucleus | |
| Tb927.3.730 | | 1.7 | | protein of unknown function, flagellar axoneme | ✓ |
| Tb927.8.3160 | | 1.7 | | protein of unknown function, cytoplasm and kDNA | |
| Tb927.11.16400 | KAP3 | 1.7 | ✓ | kinetoplast DNA-associated protein | ✓ |
| Tb927.8.2200 | | 1.7 | | terbinafine resistance locus protein (yip1), putative, endocytic | |
| Tb927.11.7600 | MC12 | 1.7 | ✓ | protein of unknown function, cytoplasm | ✓ |
| Tb927.10.9120 | | 1.7 | | protein of unknown function, interacts with a ATP-synthase-subunit protein, mitochondrion | |
| Tb927.10.8190 | | 1.6 | | T-complex protein 1, theta subunit, putative, cytoplasm | |
| Tb927.10.10330 | CDC27 | 1.6 | | Anaphase-promoting complex subunit CDC27, cytoplasm and cell tip | |
| Tb927.11.15740 | | 1.6 | | protein of unknown function, cytoplasm and flagellar axoneme | ✓ |
| Tb927.4.4630 | | 1.6 | ✓ | protein of unknown function, cytoplasm | |
| Tb927.11.10240 | HslV | 1.6 | ✓ | hslVU complex proteolytic subunit, putative | |
| Tb927.11.7370 | | 1.6 | | haloacid dehalogenase hydrolase, putative, cytoplasm and PFR | ✓ |
| Tb927.9.5280 | KRIPP16 | 1.6 | | KRIPP16, cytoplasm | |
| Tb927.11.16120 | KAP68 | 1.6 | ✓ | protein of unknown function, kDNA associated | ✓ |
| Tb927.7.6720 | | 1.5 | | protein of unknown function, only in *T. brucei*, cytoplasm | |
| Tb927.10.11060 | | 1.5 | ✓ | protein of unknown function | |
| Tb927.10.13310 | RPB5z | 1.5 | | DNA-directed RNA polymerase I subunit, putative, (RPB5z), cytoplasm and nucleus | ✓ |
| Tb927.9.3340 | | 1.5 | ✓ | protein of unknown function | |
| Tb927.11.4690 | POLIB | 1.5 | ✓ | mitochondrial DNA polymerase I protein B (POLIB), antipodal sites | |
| Tb927.10.4240 | | 1.5 | ✓ | RNA-binding protein RPC40, putative | |
| Tb927.10.15370 | RPC40 | 1.5 | ✓ | DNA-directed RNA polymerases I and III subunit RPAC1, putative, nucleus | ✓ |
| Tb927.7.6660 | J10 | 1.5 | ✓ | chaperone protein DNAj, putative | |
| Tb927.9.2310 | | 1.5 | ✓ | protein of unknown function | |
| Tb927.9.6380 | | 1.5 | | protein of unknown function, transporter family domain, endocytic | |

**Table S3. KAP68 Immunoprecipitation List**

All proteins upregulated by more than 2-fold and with a p-value ≤ 0.05 in KAP68HA-tagged versus wild-type (wt) immunoprecipitation are listed. The table includes accession number, log2 fold change, gene description, molecular weight (kDa), isoelectric point (pI), and localization according to TrypTag.org.

| gene | log2 fold change | description | kDa / pI | Tryptag |
|---|---|---|---|---|
| Tb927.11.9170 | 5.2 | DNA topoisomerase III, putative | 93kDa / 8.2 | cytoplasm;nucleus |
| Tb927.10.1900 | 3.0 | DNA topoisomerase IA, putative | 88kDa / 9.6 | antipodal site |
| Tb927.10.5830 | 2.9 | zinc finger protein, RanBP2-type, putative | 59kDa / 8.0 | endocytic, cytoplasm |
| Tb927.11.16120 | 2.9 | kDNA associated protein KAP68 | 67kDa / 10.0 | kinetoplast |
| Tb927.3.5420 | 2.8 | protein transport protein Sec24 | 100kDa / 7.1 | golgi apparatus;cytoplasm |
| Tb927.9.5020 | 2.7 | HMG-box domain containing protein mtHMG44 | 44kDa / 10.2 | kinetoplast |
| Tb927.10.6290 | 2.7 | ATP-dependent DEAD/H RNA helicase, putative | 57kDa / 9.6 | cytoplasm;nucleolus;nucleus |
| Tb927.8.3020 | 2.4 | hypothetical protein, conserved | 66kDa / 8.8 | ciliary plasm;cytoplasm |
| Tb927.7.1080 | 2.1 | hypothetical protein, conserved | 40kDa / 9.5 | cytoplasm;mitochondrion;nucleolus |
| Tb927.7.6610 | 2.1 | hypothetical protein, conserved | 96kDa / 8.9 | unknown |
| Tb927.2.1120 | 2.1 | retrotransposon hot spot protein 4 (RHS4) | | unknown |
| Tb927.11.13890 | 2.1 | AKAP7 2'5' RNA ligase-like domain containing protein, putative | 30kDa / 10.1 | cytoplasm;cytosol;mitochondrion |
| Tb927.3.1730 | 2.0 | translation elongation factor P, putative | 25kDa / 6.9 | antipodal site |
| Tb927.3.3440 | 1.8 | phytanoyl-CoA dioxygenase (PhyH), putative | 39kDa / 9.8 | cytoplasm;nucleus |
| Tb927.11.8980 | 1.6 | Elongation factor G 2, mitochondrial, putative | 90kDa / 7.2 | cytoplasm;mitochondrial matrix |
| Tb927.8.8180 | 1.6 | RNA-editing substrate binding complex protein RESC11A | 103kDa / 6.3 | cytoplasm;mitochondrion |
| Tb927.10.1450 | 1.6 | Centrin arm-associated protein 1 | 329kDa / 4.3 | microtubule structures |
| Tb927.6.2230 | 1.4 | RGG protein | 88kDa / 8.7 | kinetoplast;mitochondrion;nucleoplasm |
| Tb927.6.2100 | 1.4 | 40S ribosomal protein S30, putative | 7kDa / 12.1 | cytoplasm |
| Tb927.4.3770 | 1.4 | calcium/calmodulin-dependent protein kinase, putative | 54kDa / 8.8 | axoneme;cytoplasm;nucleus |
| Tb927.11.15640 | 1.4 | mitochondrial RNA binding complex 1 subunit | 44kDa / 5.5 | kinetoplast;mitochondrion |
| Tb927.3.2050 | 1.4 | Minicircle replication factor 172 | 172kDa / 9.5 | kinesin complex;kinetoplast;microtubule |
| Tb927.4.4160 | 1.2 | RNA-editing substrate binding complex protein RESC12 | 100kDa / 10.1 | kinetoplast;mitochondrion |
| Tb927.10.10830 | 1.2 | RNA-editing substrate binding complex protein RESC13 | 32kDa / 7.9 | cytoplasm;kinetoplast;mitochondrion;nu cleus |
| Tb927.6.4440 | 1.1 | RNA-binding protein 42 (RNA-binding motif protein 42) | 37kDa / 5.3 | cytoplasm |
| Tb927.3.1820 | 1.1 | mitochondrial RNA binding complex 1 subunit | 30kDa / 12.1 | cytoplasm;mitochondrion;nuclear lumen |
| Tb927.10.1490 | 1.1 | MKT1-like protein | 134kDa / 6.4 | cytoplasm;interchromatin granule |
| Tb927.7.880 | 1.0 | RNA-binding protein, putative | 29kDa / 5.6 | cytosol;nucleoplasm;nucleus |
| Tb927.2.3880 | 1.0 | heterogeneous nuclear ribonucleoproteins F/H homologue | 55kDa / 6.7 | cytoplasm;nucleus;ribonucleoprotein complex |
| Tb927.8.750 | 1.0 | nucleolar RNA-binding protein, putative | 39kDa / 7.9 | nucleolus;nucleoplasm;nucleus |

**Table S4. HMG44 Immunoprecipitation List**

All proteins upregulated by more than 2-fold and with a p-value ≤ 0.05 in HMG44myc-tagged versus wild-type (wt) immunoprecipitation are listed. The table includes accession number, log2 fold change, gene description, molecular weight (kDa), isoelectric point (pI), and localization according to TrypTag.org.

| gene | log2 fold change | description | kDa / pI | Tryptag |
|---|---|---|---|---|
| Tb927.11.16120 | 9.1 | kDNA associated protein KAP68 | 68kDa/ 10.0 | kinetoplast |
| Tb927.9.5020 | 8.1 | HMG-box domain containing protein mtHMG44 | 44kDa/ 10.2 | kinetoplast |
| Tb927.2.6100 | 7.8 | hypothetical protein, conserved | 53kDa/ 10.8 | mitochondrion |
| Tb927.7.2650 | 5.7 | Cytoskeleton associated protein 51V | 62kDa/ 6.0 | microtubule structures |
| Tb927.11.10780 | 5.6 | Voltage-dependent anion channel, putative | 30kDa/ 8.9 | mitochondrion OM |
| Tb927.7.5230 | 5.6 | lanosterol synthase | 103kDa/ 7.8 | cytoplasm |
| Tb927.6.3840 | 5.4 | reticulon domain protein | 21kDa/ 7.7 | microtubule structures |
| Tb927.11.15180 | 5.1 | hypothetical protein, conserved | 13kDa/ 10.8 | nucleolus |
| Tb927.9.6510 | 4.6 | Mitochondrial SSU ribosomal protein, putative | 75kDa/ 9.6 | mitochondrion |
| Tb11.1370 | 4.0 | hypothetical protein, conserved | 50kDa/ 6.1 | unknown |
| Tb927.9.4500 | 3.7 | heat shock protein, putative | 91kDa/ 9.1 | ER;mitochondrial inner membrane |

**Table S5. TAC53 Immunoprecipitation List**

All proteins upregulated by more than 2-fold with a p-value ≤ 0.05 in TAC53-HA induced versus uninduced conditions are listed. The table includes accession number, log2 fold change, gene description, molecular weight (kDa), isoelectric point (pI), and localization according to TrypTag.org.

| gene | log2 fold change | description | kDa / pI | Tryptag |
|---|---|---|---|---|
| Tb927.2.6100 | 6.2 | hypothetical protein, conserved | 53kDa / 10.8 | cytoplasm;kinetoplast;mitochondrion |
| Tb927.6.1770 | 4.3 | kinesin, putative | 69kDa / 7.1 | microtubule structures |
| Tb927.11.12750 | 4.1 | cleavage and polyadenylation specificity factor 30 kDa subunit | 31kDa / 8.3 | nucleus |
| Tb927.5.3650 | 3.9 | zinc finger protein, FYVE/PHD-type, putative | 86kDa / 10.5 | microtubule structures |
| Tb927.9.2630 | 3.0 | hypothetical protein, conserved | 208kDa / 7.7 | microtubule structures |
| Tb11.v5.0604 | 2.9 | hypothetical protein, conserved | 110kDa / 5.3 | unknown |
| Tb927.3.3940 | 2.9 | RNA-binding protein, putative | 61kDa / 6.1 | cytoplasm |
| Tb927.9.7250 | 2.7 | hypothetical protein, conserved | 20kDa / 11.2 | cytoplasm;nucleolus |
| Tb927.10.3540 | 2.5 | DNA-directed RNA polymerase I subunit RPA31 | 24kDa / 10.4 | nucleus |
| Tb927.7.4250 | 2.3 | hypothetical protein, conserved | 12kDa / 10.1 | cytoplasm |
| Tb927.10.7810 | 2.3 | Pre-rRNA-processing protein ESF1, putative | 80kDa / 4.9 | nucleolus;nucleus |
| Tb927.9.12890 | 2.3 | hypothetical protein, conserved | 16kDa / 11.5 | nucleus |
| Tb927.6.2710 | 2.3 | E3 ubiquitin-protein ligase HEL2, putative | 85kDa / 8.0 | ciliary plasm;cytoplasm |
| Tb927.7.3380 | 2.2 | Ribosomal L28e protein family | 32kDa / 4.9 | nucleus |
| Tb927.10.5670 | 2.2 | N-acetyltransferase subunit Nat1, putative | 82kDa / 7.4 | cytoplasm |
| Tb927.5.2990 | 2.1 | hypothetical protein, conserved | 27kDa / 12.4 | microtubule structures |
| Tb927.10.2810 | 2.0 | hypothetical protein, conserved | 38kDa / 4.7 | unknown |
| Tb927.9.11840 | 1.8 | pre-rRNA-processing protein PNO1, putative | 24kDa / 10.2 | cytosol;nucleolus;nucleus |
| Tb927.10.8980 | 1.8 | hypothetical protein, conserved | 17kDa / 12.3 | kinetoplast, mitochondrion |
| Tb927.3.740 | 1.8 | ZFP family member, putative | 25kDa / 8.8 | cytoplasm |
| Tb927.5.4320 | 1.7 | Pre-mRNA polyadenylation factor FIP1 | 31kDa / 6.4 | nucleus |
| Tb927.9.7670 | 1.7 | hypothetical protein, conserved | 26kDa / 10.4 | nucleus |
| Tb927.9.7110 | 1.7 | GRAM domain containing protein, putative | 32kDa / 7.8 | cytoplasm;transport vesicle |
| Tb927.11.16400 | 1.7 | kinetoplast-associated protein 3, putative | 36kDa / 12.3 | cytoplasm;kinetoplast |
| Tb927.11.8050 | 1.6 | Sas10 C-terminal domain containing protein, putative | 60kDa / 5.4 | cytoplasm;nucleus |
| Tb927.9.15290 | 1.6 | CHAT domain containing protein, putative | 149kDa / 6.3 | cytoplasm |
| Tb927.10.5300 | 1.5 | eukaryotic translation initiation factor 6 (eIF-6), putative | 26kDa / 4.8 | cytosol;nucleolus |
| Tb927.10.12490 | 1.5 | kinesin, putative | 127kDa / 7.6 | microtubule structures |
| Tb927.9.13280 | 1.5 | Double RNA binding domain protein 9 | 47kDa / 10.7 | nucleus |
| Tb927.5.4040 | 1.4 | Mitochondrial SSU ribosomal protein, putative | 94kDa / 6.0 | mitochondrion |
| Tb927.11.5230 | 1.4 | hypothetical protein, conserved | 25kDa / 6.4 | nucleus |
| Tb927.6.1470 | 1.4 | hypothetical protein, conserved | 27kDa / 12.3 | nucleus |
| Tb927.8.900 | 1.3 | splicing factor TSR1 | 37kDa / 12.2 | nucleus |
| Tb927.6.3650 | 1.3 | ADP-ribosylation factor-like protein 3C, putative | 20kDa / 7.0 | basal body |
| Tb927.3.3590 | 1.2 | U3 small nucleolar ribonucleoprotein protein MPP10, putative | 75kDa / 4.6 | nucleus |
| Tb927.8.3950 | 1.1 | hypothetical protein, conserved | 102kDa / 5.0 | nuclear pore;nucleus |

**Table S6. List of RNAi Targets**

The table includes accession number, gene description, RNAi target, position of target, and length of RNAi target.

| gene | description | RNAi target | position of target | length of target (bp) |
|---|---|---|---|---|
| Tb927.10.8980 | hypothetical protein, conserved | ORF | 4-507 | 504 |
| Tb927.1.2730 | hypothetical protein, conserved | ORF | 452-949 | 498 |
| Tb927.8.3160 | hypothetical protein, conserved | ORF | 85-486 | 402 |
| Tb927.10.13600 | Cardiolipin-dependent protein CLDP18 | ORF | 46-440 | 395 |
| Tb927.5.2970 | suppressive immunomodulating factor TSIF | ORF | 897-1395 | 449 |
| Tb927.5.2560 | hypothetical protein, conserved | 3'UTR | 35-439 | 405 |
| Tb927.10.6200 | hypothetical protein, conserved | 3'UTR | 13-305 | 293 |
| Tb927.11.1700 | hypothetical protein, conserved | ORF | 932-1399 | 468 |
| Tb927.7.7090 | hypothetical protein, conserved | ORF | 82-436 | 355 |
| Tb927.5.2630 | hypothetical protein, conserved | ORF | 113-606 | 494 |
| Tb927.2.6100 | TAC53 | ORF | 95-426 | 332 |
| Tb927.2.6100 | TAC53 | 3'UTR | 38-459 | 422 |
